# Developing Bioactive Hydrogels Containing Cell-derived Extracellular Matrix: Implications in Drug and Cell-free Bone and Cartilage Repair

**DOI:** 10.1101/2024.03.04.583366

**Authors:** Ali Coyle, Aishik Chakraborty, Jiaqi Huang, Yasmeen Shamiya, Wei Luo, Arghya Paul

**Affiliations:** School of Biomedical Engineering, The University of Western Ontario, London, ON N6A 5B9, Canada; Department of Chemical and Biochemical Engineering, The University of Western Ontario, London, ON N6A 5B9, Canada; Collaborative Specialization in Musculoskeletal Health Research and Bone and Joint Institute, The University of Western Ontario, London, ON N6A 5B9, Canada; Department of Chemistry, The University of Western Ontario, London, ON N6A 5B9, Canada

**Keywords:** cell-derived extracellular matrix (ECM), 3D printing, bone regeneration, cartilage regeneration, mesenchymal stem cells

## Abstract

The prevalence of osteoarthritis has been increasing in aging populations, which has necessitated the use of advanced biomedical treatments. These involve grafts or delivering drug molecules entrapped in scaffolds. However, such treatments often show suboptimal therapeutic effects due to poor half-life and off-target effects of drug molecules. This study aims to overcome these limitations by 3D printing gelatin-based hydrogel scaffolds containing cell-derived extracellular matrix (ECM) as the bioactive therapeutic cargo. Here, pre-osteoblastic and pre-chondrogenic murine cells were differentiated *in vitro*, decellularized, and incorporated into methacrylated gelatin (GelMA) solutions to form osteogenic (GelO) and chondrogenic (GelC) hydrogels, respectively. The integration of the bioactive decellularized extracellular matrix (dECM) allows GelO and GelC to induce differentiation in human adipose-derived stem cells (hADSCs) toward osteogenic and chondrogenic lineages. GelO and GelC can be covalently adhered using carbodiimide coupling reaction, forming bioactive osteochondral plug. Moreover, this osteochondral plug can also induce differentiation of hADSCs. To conclude, this ECM-based bioactive hydrogel offers a promising new drug-free and cell-free treatment strategy for bone and cartilage repair, and future osteoarthritis management.

## 1. Introduction

Osteoarthritis (OA) and osteochondral defects (OCDs) are debilitating joint disorders characterized by progressive cartilage degradation and underlying bone abnormalities[1,2]. These conditions affect millions of individuals globally, leading to pain, loss of function, and reduced quality of life. Therefore, it is crucial to understand the prevalence of OA and OCDs and the limitations of current treatment modalities to explore novel therapeutic approaches. The current gold standard treatment options for OA and OCDs primarily aim to manage pain, relieve symptoms, or replace damaged tissue with transplantation and implantation[3]. These approaches include grafting techniques, such as xenografts, allografts, and autografts. However, they pose several challenges, including (a) immune rejection, (b) the potential transmission of zoonotic diseases are major concerns associated with xenografts[4], (c) variable long-term outcomes[5], and (d) limited availability of suitable autograft donor sites. Countering these challenges, synthetic biomimicking alloplastic materials, such as bioglass, hydroxyapatite, and collagen sponge have been developed, showing promising therapeutic outcome. However, challenges remain in achieving proper integration with native tissue, long-term stability[6], natural remodelling capabilities[7], and limited control over structure[8].

Recent advancements in therapies involving the use of cells, proteins, and peptides either alone or in combination have revolutionized the field of OA and OCDs treatment. One of the most promising cell-based therapies for osteochondral repair involves the use of stem cells[9]. However, the heterogeneity and variability of MSCs obtained from different sources and donors present a significant limitation in stem cell-based therapies for OA[10]. In addition to cell-based therapies, growth factors have emerged as essential components in the regeneration of damaged osteochondral tissue. Transforming growth factor-beta (TGF-β) and insulin-like growth factor-1 (IGF-1) are two growth factors that have shown promising results in promoting cartilage formation and enhancing tissue healing[11]. But aside from their high cost, there is a risk of off-target effects and excessive tissue growth when growth factors are administered in high doses[12]. Peptides have also gained attention as potential therapeutic agents for osteochondral repair, but display poor stability[13] and bioavailability. As a result, it is necessary to design therapeutic strategies that can limit the use of biobased drugs or cells.

ECM-based therapeutics have become promising alternatives to biobased drugs or cells for the treatment of OA and OCD. ECM can be harvested from various organs and tissues which involves the removal from living organisms or cadavers, followed by decellularization. In the field of osteochondral repair, where the regeneration of both cartilage and underlying bone is crucial, ECM extracted from cartilage and bone tissues has demonstrated significant potential[14]. While tissue-derived ECM offers several advantages in terms of obtaining a native ECM composition and structure, it is not without limitations. One major drawback is the limited availability of donor organs or tissues, especially for specific applications[15]. Besides availability, tissue-derived ECM also suffer from high batch-to-batch variability. As a replacement, ECM-derived from two-dimensional tissue cultures has been proposed[16]. By culturing cells in a controlled laboratory environment, cell-derived ECM offers the advantage of scalability and consistency.

Here, we have developed water-swellable three dimensional (3D), porous polymeric hydrogel scaffolds containing dECM from murine pre-osteoblastic and pre-chondrogenic cell lines. The cells were first differentiated for fourteen days to develop a nutrition-rich ECM. Subsequently, the culture was decellularized, and the extracted dECM was added to GelMA, which is a versatile biomaterial derived from gelatin. GelMA possesses several advantages, including biocompatibility, tunable mechanical properties, and excellent cell adhesion, making this biopolymer as our primary choice for the hydrogel scaffold[17]. The mixture of GelMA along with the dECM were then 3D printed using a digital light processing (DLP)-based bioprinter to obtain osteogenic, GelO and chondrogenic, GelC scaffolds. Finally, we could adhere the two hydrogels to form osteochondral plugs with the potential of tissue repair. **Figure 1** describes the steps involved with the fabrication of the osteochondral plug. Overall, this research contributes to the advancement of tissue engineering approaches for osteochondral tissue repair and brings us closer to providing effective and minimally invasive treatments for patients suffering from OA and OCDs.

**Figure 1.**
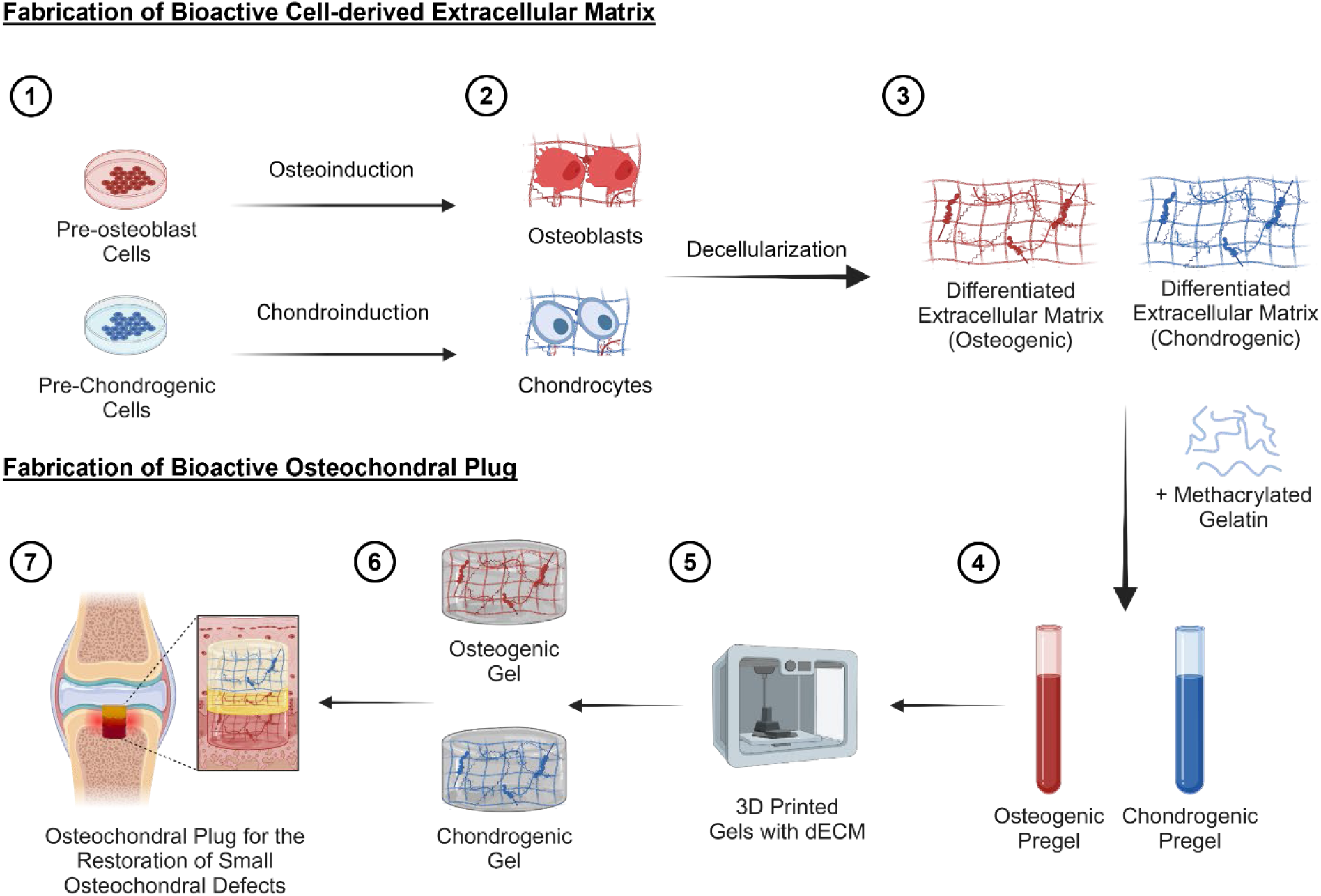
Graphical representation of osteogenic and chondrogenic GelMA hydrogel containing dECM for osteochondral repair. Pre-osteoblast cells and pre-chondrogenic cells were first cultured in a controlled laboratory environment (1). The cells were then differentiated toward osteogenic and chondrogenic lineages for 14 days (2). The deposited extracellular matrix was extracted by decellularization (3). The differentiated extracellular matrices were added to methacrylated gelatin to prepare osteogenic and chondrogenic pregels (4). The pregels were then 3D printed using a light-based 3D bioprinter. (5) Subsequently, gels capable of inducing osteogenesis and chondrogenesis in stem cells were prepared and characterized (6). Finally, the gels were adhered using carbodiimide coupling reaction to form osteochondral plug suitable for repairing small osteochondral defects.

## 2. Materials and Methods

### 2.1. dECM extraction from pre-osteoblastic murine cells (MC3T3-E1)

Pre-osteoblastic MC3T3-E1-E1 cell line (ATCC, CRL-2593, subclone 4) were grown in an expansion medium which contained α-Minimum Essential powdered Medium (α-MEM; Sigma, M0644) dissolved in autoclaved DI water and supplemented with 10% Fetal Bovine Serum (FBS; Sigma, F1051) and 1% PenStrep (PS; Gibco, 15070-63). For passaging, cells were treated with 0.25% Trypsin-EDTA (Gibco, 25200072) for 10 minutes in an incubator (Thermo Forma Steri Cycle 370, US). Passaging was performed at 90% cellular confluency. To induce osteogenesis in the MC3T3-E1 cell monolayer, 5mL of Osteogenic Differentiating Media (ODM) was used when cells reached confluency with a total change of media every 3 days. ODM was made using the basal medium supplemented with 10mM β-glycerophosphate disodium salt hydrate (BGP, sigma, G9422) and100 nM Dexamethasone (Dex, Sigma, D4902). For ECM production, 1.5×10^6^ Cells were seeded onto 100 mm Petri dishes and incubated at 37 °C with 5% CO_2_. After 14 days of differentiation without cell splitting, the ECM monolayer was treated with 1% Triton X-100 in PBS (v/v) with 1% PS for 10 minutes commonly used in decellularization studies[18]. The differentiated dECM monolayer (OdECM) was washed 4 times with 5 mL of PBS with 1% PS and then washed in 5mL of water overnight at 4 °C. After the removal of excessive water, the monolayer was almost airdried in a biosafety cabinet. Damp dECM was scrapped from the petri dish, lyophilized, and stored at -80°C until use.

### 2.2 dECM extraction from pre-chondrogenic murine cells (ATDC5)

The pre-chondrogenic ATDC5 cell line was a gift from Dr. Frank Beier (Western University, Canada). The cells were cultured in expansion medium Dulbecco’s modified Eagle’s medium with nutrient mixture F-12 (DMEM/F12; Thermo Fisher, 11320033) and supplemented with 5% FBS, 1% PS, 10mM BGP, and 0.5mM L-ascorbic acid (Sigma, A4544). For passaging, cells were trypsinized for 5 minutes in the incubator. To induce chondrogenesis in the ATDC5 cell monolayer, 5mL of Chondrogenic Differentiating Media (CDM) was used with a total change of media every 3 days. CDM was made using the expansion medium supplemented with 1% ITS 100X (Sigma, I3146). For ECM production, 1.5×10^6^ Cells were seeded onto 100 mm Petri dishes and incubated at 37 °C with 5% CO_2_. After 14 days of differentiation without cell splitting, the ECM monolayer was treated with 1% Triton X-100 in PBS (v/v) with 1% PS for 10 minutes. The dECM monolayer (CdECM) was washed 4 times with 5 mL of PBS with 1% PS and then washed in 5mL of water overnight at 4 °C. After the removal of excessive water, the monolayer was almost airdried in a biosafety cabinet. Damp dECM was scrapped from the petri dish, lyophilized, and stored at -80°C until use.

### 2.3. *In vitro* feasibility analysis using human adipose-derived stem cells (hADSCs)

The hADSCs used in the experiment were obtained from Lonza (Cat. No. PT-5006). These cells were cultured in ADSC basal medium (Lonza, Cat. No. PT-3273), which was supplemented with the ADSC-GM SinglequotsTM Supplement Kit (Lonza, Cat. No. PT-4503). Cells were incubated at 37 °C at 5% CO2.

For dECM optimization, confluent cultures were treated once with 0.4% (w/v) OdECM and 0.3% CdECM, respectively, in culture media with media change every 3 days. To test for osteogenesis and chondrogenesis of GelO and GelC, 5×10^4^ were seeded in 24-well plates at which the bottom of the wells was covered with a thin layer of GelO, GelC or both.

For positive controls, Dulbecco′s Modified Eagle′s Medium (DMEM, Sigma, D5030) was used and supplemented with 10% FBS, 1% PS, 0.5mM L-ascorbic acid, and 10mM BGF. To make an osteoinductive (OI) medium, 100 nM Dex was added and for a chondroinductive (CI) medium 100 nM Dex and 1% ITS were added to the media. All experimentations involving hADSCs were carried out for 21 days.

### 2.4 Scanning Electron Microscopy and Energy-dispersive X-ray Spectroscopy

The dECM and bioactive hydrogel samples underwent analysis using SEM/EDX following an established protocol[19], which combines scanning electron microscopy and energy-dispersive X-ray spectroscopy. A Hitachi SU3500 variable pressure SEM and an Oxford X-Max50 SDD X-ray detector were employed for this purpose. SEM generated surface topography images, while EDX, a semi-quantitative technique, could detect elements ranging from carbon to uranium. Its minimum detection limit was approximately 0.1 weight %, and it probed the sample to a depth of a few micrometres. The SEM/EDX analyses were conducted at an accelerating voltage of 20.0 kV. To minimize artifacts caused by sample charging, a thin layer of gold coating was applied to the samples.

### 2.5 Cytocompatibility of the hydrogels using MTS assay

The hADSCs were seeded at a density of 15000 cells per well in 96-well plates and cultured overnight. In this study, two cytocompatibility tests were performed using cell proliferation MTS assay (Promega, Cat. No. G3580). This assay involved the conversion of MTS [3-(4,5-dimethyl-2-yl)-5-(3-carboxymethoxyphenyl)-2-(4-sulfophenyl)-2H-tetrazolium] into formazan by living cells in a reducing environment. In the first experiment, the cells were exposed to different concentrations of CdECM and OdECM suspended in stem cell media (0.1, 0.3, 0.4, 0.5, and 0.6 (w/v)) for durations of 24 h. In the second experiment, thin layers of GelO and GelC were photo-crosslinked with blue light (470 nm) on the bottom of 96-well plates. They were washed with DI water and PBS. The same number of hADSCs were seeded on coated wells and cultured overnight. After 24 hours, 10 μL MTS reagent was added to each well of the 96-well plate and incubated for 4 h in darkness. After incubation, the formazan supernatant was transferred to a fresh plate, and the absorption values were measured at a wavelength of 490 nm using a microplate reader.

### 2.6 Quantitative Proteomics of OdECM and CdECM to identify their protein composition

A total phenotype of OdECM and CdECM was characterized by quantitative proteomic analysis using the Orbitrap Fusion^TM^ Lumos^TM^ mass spectrometer. Functional enrichment analysis of identified proteins were conducted with the bioinformatics platform STRING database (http://string-db.org) and enriched gene oncology terms under biological processes, cellular components and molecular functions were visualized using bioinformatics platform (https://www.bioinformatics.com).

### 2.7 Preparation and characterization of hydrogels

GelMA was prepared using a previously established method[20]. Initially, Gelatin from porcine skin (Type A) (Sigma, G2500) was dissolved in 10% PBS (w/v) at 60°C under constant stirring for 1 hour. After the complete dissolution of the gelatin, methacrylic anhydride (Sigma, 276685) (0.8 mL/g of gelatin) was slowly added dropwise to the gelatin solution while maintaining an alkaline pH. The reaction mixture was left to rotate for an additional 2 hours at the same temperature. Subsequently, the GelMA solution was diluted with heated 100 mL PBS and subjected to dialysis for approximately 1 week using a dialysis membrane with a molecular weight cut-off of 12-14 kDa and distilled water maintained at 50°C. The water used in the dialysis process was changed twice daily to remove any unreacted methacrylic anhydride, ensuring the elimination of potential toxicity. The resulting GelMA solution was then lyophilized to obtain a powdered form. By varying the molar ratio of methacrylic anhydride to gelatin, the degree of methacrylation was adjusted to about 73%.

### 2.8 Degradation of hydrogels

To assess the impact of dECM particles on the degradation behaviour, GelMA, GelO, and GelC hydrogels were prepared with a volume of 150 µL and lyophilized. The experiment involved collecting samples with 3 replicates on day 10. The hydrogels were placed in 24-well plates and immersed in 2 mL of PBS containing 1% PS. The plates were placed in a shaker incubator with a temperature of 37.0°C and a speed of 30 RPM. The hydrogels were freeze-dried and weighed at the specified time point. The degradation rate (%) was determined using the following equation:

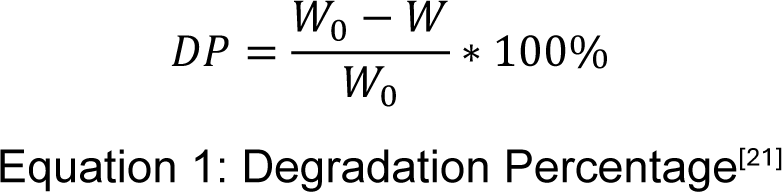

Here, W_0_ represents the initial weight of the dried hydrogel, W corresponds to the weight of the dried hydrogel at day 10, and DP is the degradation percentage.

### 2.9 Swelling of hydrogels

Swelling ratio of the hydrogels were determined following established protocol[19].To evaluate the impact of dECM particles on the swelling characteristics of hydrogels, GelMA, GelO, and GelC hydrogels were prepared (n = 5) with a volume of 150 µL each. The dried weights of the hydrogels were recorded as W_0_. Then, dried hydrogels were placed into separate wells of a 24-well plate containing 2 mL of PBS at a temperature of 37.0°C. At intervals of 30 minutes for 7 hours, the mass of each hydrogel was measured to calculate the swelling ratio using the following equation:

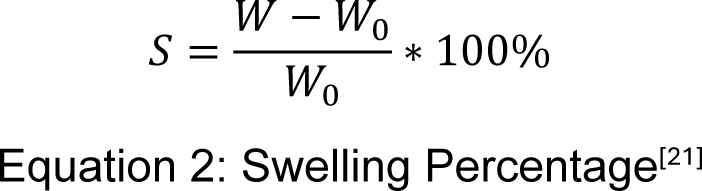

where S represents the swelling ratio expressed as a percentage, W represents the weight of the swollen gel, and W_0_ denotes the initial dry weight.

### 2.10 Compressive modulus of hydrogels

To determine the compressive moduli of hydrogels, mechanical testing was conducted using INSTRON Force Transducer Model 2519-101 under unconfined uniaxial conditions[22]. Three groups of cylindrical hydrogels were 3D-printed using DLP. The groups with 5 replicates were GelMA, GelO, and GelC with a diameter of 3mm and height of 6 mm. To prevent slippage during the test, sandpaper was applied to the compression probes. The testing was performed on the swollen hydrogels at 0.3 mm/min compression rate, and the compressive modulus was determined by analyzing the linear portion of the stress-strain curve within the 1-15% strain range. The samples were compressed until fracture occurred. The compressive modulus profile was determined using *Equation 3*, in which 𝜎𝜎 represents the applied compressive stress, and 𝜀𝜀 represents the strain.

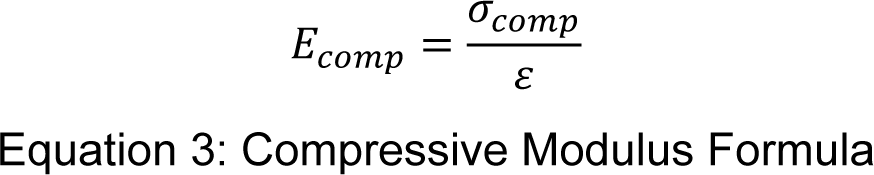

### 2.11 Rheological analysis of hydrogels

The rheological studies were conducted using a HAAKE Modular Advanced Rheometer System (MARS) equipped with a P20/Ti titanium plate as the measuring geometry provided by Fischer Scientific, USA. The experiments were performed at 37 °C to simulate physiological conditions. Cylindrical GelMA, GelO, and GelC samples with dimensions of 6 mm × 6 mm were printed with three replicates. Before the rheological measurements, the hydrogels were hydrated in PBS at pH 7.4 for 5 minutes to prevent stickiness during testing. Any excess liquid on the gel surfaces was gently dabbed off to maintain uniformity. The rheological properties were evaluated through stress sweeps ranging from 0.1 to 104 Pa, applying different levels of stress to the hydrogel samples and measuring their corresponding strain responses.

### 2.12 Carbodiimide-based coupling reaction for adhering Gel-O to Gel-C

To chemically bond GelO and GelC hydrogels, an adhesive solution was synthesized using an established protocol[23]. To prepare the adhesive solution, 2-(N-morpholino)ethanesulfonic acid (MES) sodium salt was dissolved in DI water at a concentration of 0.1 mol/L at room temperature, resulting in a 0.1 M MES buffer solution. The pH of the MES buffer solution was adjusted to 6 by gradually adding hydrochloric acid solution. Gelatin at a concentration of 20% (w/v) was dissolved in Eppendorf tubes using the MES buffer, and the tubes were then placed in a 60°C environment for 10 minutes to aid in the dissolution process. N-(3-Dimethylaminopropyl)-N’-ethylcarbodimide hydrochloride (EDC, Sigma, E7750-1G) at a concentration of 10% (w/v) was dissolved in Eppendorf tubes using the 0.1 M MES buffer at room temperature. Similarly, N-Hydroxysuccinimide (NHS, Sigma, 130672-5G) at a concentration of 5% (w/v) was dissolved in Eppendorf tubes using the 0.1 M. The adhesive components were mixed right before applying between the two hydrogels and left to rest at room temperature for 1 hour.

### 2.13 Alizarin red S staining (ARS) to demonstrate osteogenesis

The ARS method was employed to assess mineralization in cell culture samples and hydrogel constructs based on the manufacturer’s protocol [24]. Initially, confluent MC3T3-E1 cells were cultured with growth media or ODM for 14 days. The monolayers were washed with PBS and fixed using a 4% paraformaldehyde solution to preserve the cellular and extracellular matrix components. Subsequently, the fixed samples underwent rinsing with diH_2_O to eliminate any residual fixative. To visualize the mineralized areas, the samples were stained with 40 mM Alizarin Red (ScienCell, Cat. No. 8678) at room temperature, which selectively binds to calcium deposits. After 30 minutes, the samples were thoroughly washed to remove excess dye and reduce background interference. The stained samples were then examined under a microscope to visualize the mineralized regions, indicated by the red staining. Quantification of mineralization was performed through colorimetric analysis by extracting the dye and measuring the absorbance at 405nm wavelength. To ensure accurate quantification, calibration curves using known concentrations of calcium standards were utilized. Similarly, ARS was performed on MSCs that were grown on GelO for 21 days.

### 2.14 Alcian Blue Staining to demonstrate chondrogenesis

Confluent ATDC5 cells were cultured with growth media or CDM for 14 days. The monolayers were washed with PBS and fixed using 95% methanol for 20 minutes. A solution of 1% Alcian Blue 8GX (Thermo Fisher, J60122) in 0.1 M HCl was prepared. After washing off the methanol, the monolayers were exposed to the dye solution overnight. Following this, the samples were thoroughly washed with PBS and then diH_2_O. Images of the stained cells were captured. Similarly, ABS was performed on MSCs that were grown on GelC for 21 days.

### 2.15 Sulfated-Glycosaminoglycans (S-GAG) Quantification to prove chondrogenesis

To evaluate chondrogenesis by GelC, hADSCs cultured on a thin layer of hydrogel for 21 days were subjected to Dimethylmethylene Blue Assay (DMMB, ABCAM, ab289846) based on the manufacturer protocol. Briefly, 100 mg of Cell/hydrogel samples were transferred to Eppendorf tubes, followed by the addition of ice-cold homogenization buffer. The resulting mixture was centrifuged at 12000 x g and 4°C for 20 minutes. The supernatant, obtained after centrifugation, was collected for subsequent analysis. In a 96-well plate, 50 µL of the supernatant was added to a well labelled as “Sample,” while another well labelled as “Sample Control” received 50 μL of homogenization buffer. Both wells were adjusted to 100 μL using S-GAG Assay Buffer. To create the standard curve, a diluted s-GAG Standard solution was prepared by combining s-GAG stock Standard with s-GAG Assay Buffer. Subsequently, 200 μL of s-GAG Dye was added to all wells. After a 2-minute incubation at room temperature, the absorbance of all wells was measured at 525 nm.

### 2.16 DAPI/Phalloidin Staining to determine cell morphology

DAPI/phalloidin staining technique was employed to visualize nuclear DNA and filamentous actin. Samples were washed with PBS followed by the addition of 4% paraformaldehyde for a 5-minute incubation at room temperature. Cells were then treated with 0.1% Triton X and incubated at room temperature for 10 minutes. After three washes with PBS, a 1:40 dilution of phalloidin was added to the samples and incubated at room temperature for 30 minutes. Following another round of three washes with PBS, a 1:1000 dilution of DAPI (Thermo Fisher, D1306) was added and incubated at room temperature for 3 minutes. Subsequently, the samples were washed with PBS and mounted. The mounted samples were subjected to fluorescence microscopy to examine the morphology of actin filaments, which are pivotal in regulating cell shape, polarity, and crucially, cell motility.

To assess the successful decellularization of differentiated ECM, DAPI staining was performed on CdECM and OdECM after 14 days.

### 2.17 Reverse Transcription-Quantitative Polymerase Chain Reaction (RT-qPCR) Analysis to quantify gene-markers associated with osteogenesis and chondrogenesis

RT-qPCR analysis was carried out for hADSCs grown on 24-well plates and treated with 0.3% CdECM, 0.4% OdECM, GelO, GelC, or GelO-GelC for 21 days. RNA isolation was performed using the RNeasy mini kit (Qiagen, Cat. No. 74104) according to the manufacturer’s protocol. Briefly, cells were harvested from the monolayer and lysed using 350 µL of lysis buffer, ensuring the release of RNA while preserving its integrity. The lysate was then mixed with 70% ethanol to facilitate RNA binding to the silica membrane of the RNeasy spin column. The column was subsequently washed to remove impurities, and the bound RNA was eluted in a separate collection tube. The extracted RNA was quantified and assessed for quality using NanoDrop (Thermo Scientific, Canada). Five hundred nanograms of total RNA were used to synthesize cDNA using a High-Capacity cDNA Reverse Transcription Kit (Thermo Fisher, Cat. No. 4368814). Reverse transcription was performed in a Chromo4TM Real-Time PCR Thermocycler (BioRad, Canada).

TB Green Advantage qPCR premix (Takara Bio, Cat. No. 639676) was used for the amplification of osteogenic and chondrogenic markers. The PCR reaction mixture was prepared by combining the TB Green Advantage qPCR Premix, PCR forward and reverse primers (See Table 1 for primer sequences), template cDNA, and nuclease-free water according to the recommended volumes and concentrations provided. The target osteogenic markers were Runt-Related Transcription Factor 2 (*RUNX2)*, osteocalcin (*OCN*), and osteopontin (*OPN*); the chondrogenic markers were aggrecan (*ACAN*), SRY-Box Transcription Factor 9 (*SOX9*), and Matrix metalloproteinase 13 (*MMP13*); and the reference gene was *GAPDH* (housekeeping gene). After gentle centrifugation, the samples were placed in LightCycler® 96 System (Roche, Canada). The reaction was initiated with an initial denaturation step at 95°C for 10-30 seconds with detection turned off. Subsequently, combined annealing and extension were carried out at a temperature range of 55°C to 72°C for 15 seconds with detection turned on. The cycle number was set to 45 cycles. Following the completion of the reaction, the amplification curve and melting curve were analyzed for further assessment. After establishing a standard curve for each primer set, quantification of gene expression was performed using Pfaffl Method[25] with *GAPDH* and control groups being the reference gene and the calibrator, respectively.

**Table 1.**
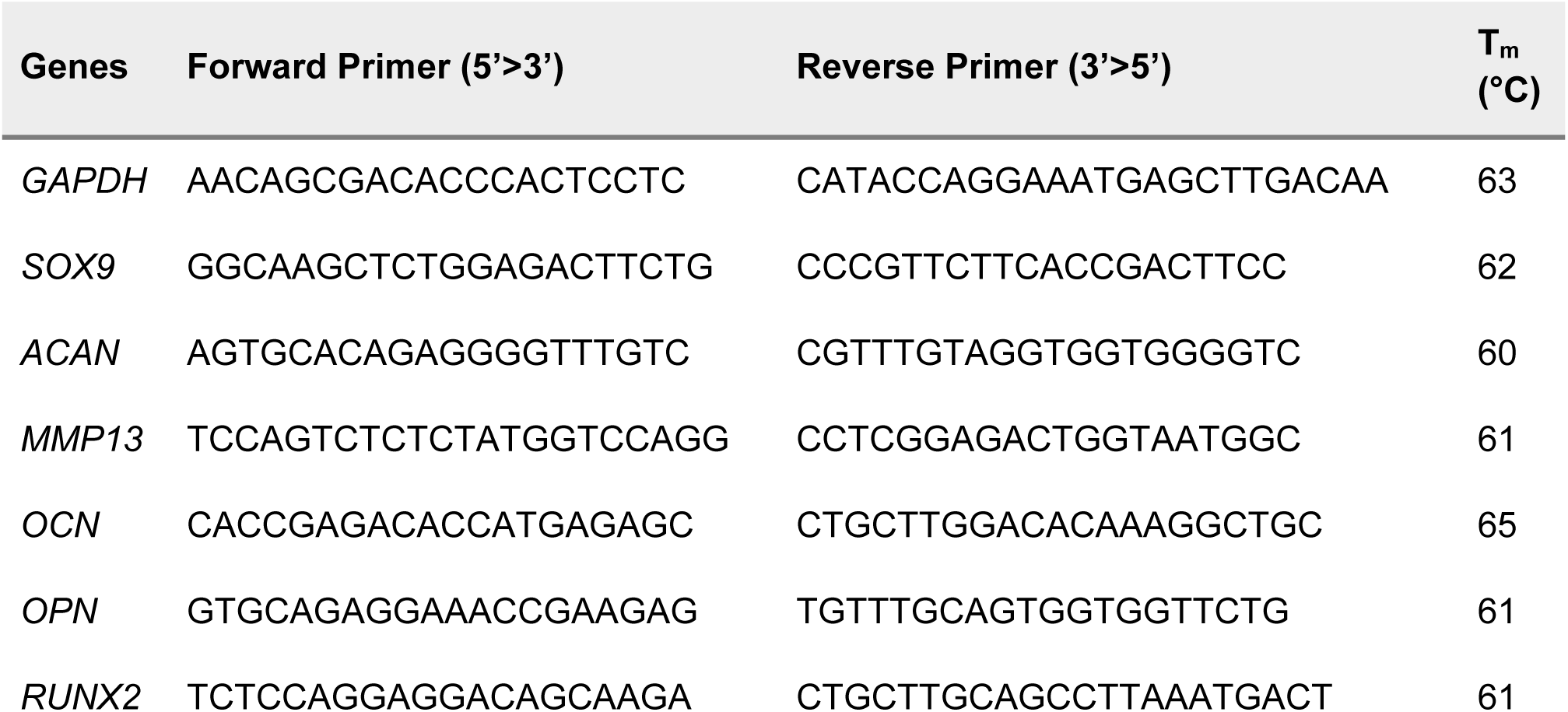
RT-qPCR forward and reverse primer sequences

### 2.18 Statistical analysis

The experimental data were expressed as the mean ± standard deviation. Differences between the experimental groups were assessed using statistical tests appropriate for the group size: the Student’s t-test for two groups, one-way ANOVA for three or more groups, followed by Tukey post hoc comparisons. Statistical significance was defined as a p-value less than 0.05. (* = p<0.05, ** = p<0.01, *** = p<0.001, **** = p<0.0001).

## 3. Results and Discussion

### 3.1 Production of dECM from differentiated MC3T3-E1 and ATDC5 cell lines

Firstly, MC3T3-E1 and ATDC5 cells were grown to confluency on 100 mm Petri dishes and differentiated using osteogenic differentiating media (ODM) and chondrogenic differentiating media (CDM), respectively, for 14 days. MC3T3-E1 is a pre-osteoblastic cell line derived from calvarial tissue of newborn mice that have been extensively utilized to study osteoblast differentiation and bone formation[26]. ATDC5 is a mouse pre-chondrogenic cell line derived from teratocarcinoma, which possesses characteristics of both chondrocytes (cartilage cells) and pre-adipocytes[27]. ATDC5 cells are widely used in research focused on cartilage development and chondrogenesis. Compared to other cells such as stem cells, the advantages of using these cell lines are but not limited to availability and ease of use, excessive ECM synthesis in a short time, excellent representation models, reproducibility, compatibility with molecular techniques, and standardized differentiation protocols[26–28].

Supported by previous works[29,30], dexamethasone served as the primary agent to induce osteogenesis in ODM, while for the preparation of CDM, a combination of human insulin, transferrin, and selenium (ITS) was utilized for chondroinduction. Furthermore, to enhance ECM deposition, L-ascorbic acid and BGP were used in the preparation of the differentiating media. L-ascorbic acid is a powerful antioxidant that plays a crucial role in the synthesis of collagen, a major component of the ECM matrix and enhances cell proliferation[31]. BGP, on the other hand, is an organic phosphate compound that can induce mineralization in osteoblasts, and it has been shown to aid in the differentiation of osteoblasts and chondrocytes[32,33]. These two reagents are commonly used in culture systems aiming to study bone or cartilage development.

Differentiated osteoblastic and chondrogenic cell cultures were then treated with 1% Triton-x100 to produce OdECM and CdECM. Triton-X100 effectively solubilizes cell membranes, disrupting cellular structures and facilitating the removal of cellular contents with the preservation of the inherent bioactive molecules[34]. The OdECM was scarped from the culture dish, washed, and then lyophilized to be used for further experiments. Each plate of MC3T3-E1 produced 2.22 ± 0.40 mg of OdECM, and each plate of ATDC5 produced 2.80 ± 0.72 mg CdECM in their lyophilized form. Overall, this technique utilizes commonly accessible laboratory equipment and resources, making it convenient to implement within a typical laboratory environment.

### 3.2 Differentiated MC3T3-E1 cell-derived OdECM is capable of inducing osteogenesis in hADSCs

We first characterized and evaluated the OdECM extracted from differentiated MC3T3-E1 cells. Figure 2A demonstrates the successful removal of cellular material using DAPI staining, indicating the effectiveness of the decellularization process. This finding aligns with previous studies that have utilized Tritox-100 to remove cellular components from cell culture and generate acellular scaffolds[34]. SEM images in Figure 2B reveals structural differences such as mineralization between dECM derived from differentiated (+) and non-differentiated (-) MC3T3-E1 cells, respectively. These differences in dECM architecture may be attributed to the influence of cellular differentiation on the organization and composition of the dECM. Previous studies have reported that cell differentiation can alter the expression of ECM components, leading to variations in dECM structure[35]. Energy dispersive x-ray spectral chart in Figure 2C shows relatively higher amounts of mineral deposition such as calcium (Ca^2+^) and magnesium (Mg^2+^) in the OdECM samples. The mineralization potential of OdECM is particularly significant for bone tissue engineering applications, as the presence of Ca^2+^ is an indication of osteogenesis[36]. The detection of minerals in the differentiated dECMs suggests that they possess the necessary components to support bone regeneration and is in line with previous studies that have reported successful mineralization of dECM derived from various cell types[37]. The Alizarin Red staining in Figure 2D confirms the mineralization potential of the differentiated OdECM. As discussed, the deposition of calcium in the matrix indicates its ability to support mineralization processes. Figure 2E quantifies the relative calcium levels in the groups through dye extraction from ARS. The higher calcium levels in the differentiated dECM group suggest its superior mineralization capacity compared to the non-differentiated group. The positive staining and higher calcium deposition in the differentiated dECM group further support its potential as a scaffold for bone tissue engineering.

**Figure 2.**
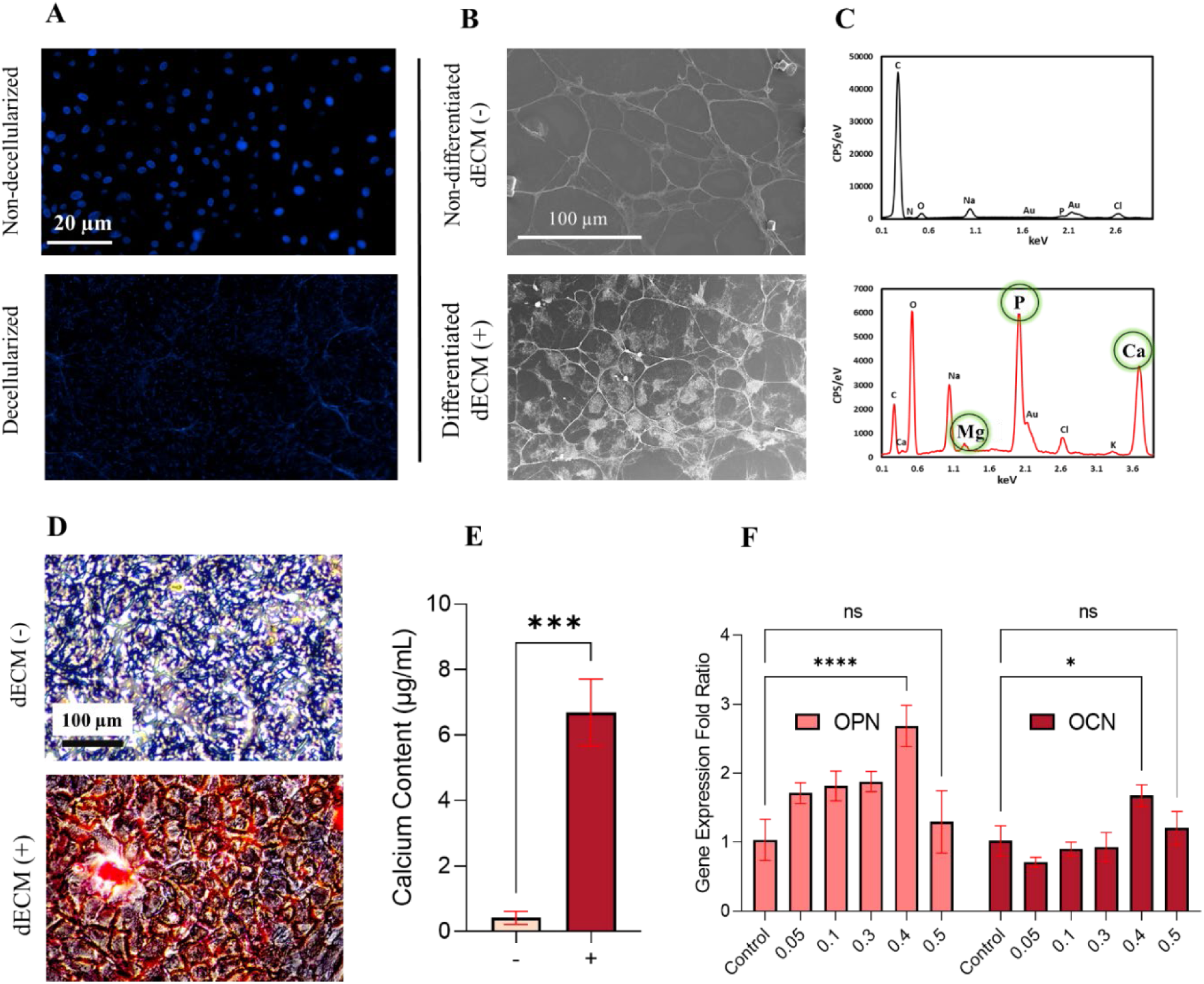
Characterization of OdECM obtained from MC3T3-E1 cell line. A) DAPI staining of MC3T3-E1 cells before (up) and after (down) decellularization, indicating successful removal of cellular material. B) Scanning Electron Microscopy (SEM) images comparing dECM derived from differentiated (+) and non-differentiated (-) MC3T3-E1 cells, showing structural differences. C) Energy-dispersive X-ray (EDX) analysis of the two groups, revealing relatively higher amount of minerals (Ca^2+^, Mg^2+^) in the differentiated dECM samples. D) Alizarin Red staining of the MC3T3-E1 cells, confirming the mineralization potential of the differentiated dECM. E) Quantification of dye extraction from Alizarin Red staining, demonstrating the relative calcium levels in the groups (n=3). F) Reverse Transcription-quantitative Polymerase Chain Reaction (RT-qPCR) analysis of osteogenic markers (*OPN* and *OCN*) in human adipose-derived stem cells (hADSCs) treated with varying concentrations of OdECM (w/v), normalized to the *GAPDH* reference gene (n=3). 0.4% (w/v) concentration showed the highest gene expression (* p<0.05, *** p<0.001, **** p<0.0001).

In Figure 2F, RT-qPCR analysis investigates the expression of osteogenic markers (*OPN* and *OCN*) in hADSCs treated with varying concentrations of OdECM. *OPN* gene (also known as *SPP1*) encodes the protein osteopontin, which is involved in various functions related to bone remodelling, including cell adhesion, matrix mineralization, and regulation of osteoblast activity[38]. It contributes to the regulation of bone formation, repair, and remodelling processes. *OCN* gene (also known as *BGLAP*) encodes the protein osteocalcin, which is a non-collagenous protein found in bone and dentin[39]. Osteocalcin plays a crucial role in regulating mineralization and the maturation of the ECM during bone formation. It is considered a marker of mature osteoblasts and is involved in the organization and deposition of hydroxyapatite crystals, contributing to bone mineralization[39]. These genes are vital components of the complex regulatory network that governs osteogenesis and the maintenance of skeletal health.

From the RT-qPCR analysis, it was found that a concentration of 0.4% (w/v) of OdECM leads to the highest expression of osteogenic markers. For this reason, a concentration of 0.4% was used for all other experimentations. This finding aligns with a related study that demonstrated successful bone formation in a mouse model using a PEG hydrogel incorporated with partially digested dECM derived from MC3T3-E1 cells, with a concentration of 0.48% (w/v)[40]. To understand the underlying mechanism behind the expression of osteogenic markers (*OPN* and *OCN*), several factors come into play. OdECM likely provides a rich source of signalling molecules, such as growth factors, cytokines, and matrix proteins, which interact with cell surface receptors and trigger intracellular signalling pathways. One such pathway is the transforming growth factor-beta (TGF-β) signalling pathway, known to play a critical role in osteogenesis[41]. TGF-β stimulates the expression of osteogenic markers through the activation of downstream effectors such as Smads and Runx2[41]. These effectors regulate the transcriptional activity of osteogenic genes, leading to enhanced expression of *OPN* and *OCN*. These results provide valuable insights into the characterization of OdECM derived from differentiated pre-osteoblastic cells. The successful decellularization, distinct structural differences, mineralization potential, and induction of osteogenic markers in hADSCs support the suitability of differentiated dECM for bone tissue engineering applications.

### 3.3 Differentiated ATDC5 cell-derived CdECM is capable of inducing Chondrogenesis in hADSCs

Next, we characterized and evaluated the CdECM extracted from differentiated ATDC5 cells, as shown in Figure 3. DAPI images shown in Figure 3A capture the removal of DNA, suggesting successful decellularization. Figure 3B displays SEM images revealing the structural changes between dECM extracted from differentiated and non-differentiated ATDC5 cells. The EDX spectra in Figure 3C demonstrates the increased production of inorganic components, including phosphates (P) and calcium (Ca^2+^) in the CdECM samples. Phosphate plays a crucial role in cartilage mineralization, cell differentiation, and apoptosis and has been implicated as a rate-limiting factor in the process of cartilage repair[42]. Through experiments using ATDC5 cells, it was found that phosphate promoted matrix mineralization and the expression of collagen X, a marker of chondrocyte maturation which highlights phosphate’s regulatory role in chondrocyte maturation and apoptosis[43]. Alcian Blue staining was performed on the ATDC5 cells to visualize the presence of sulfated glycosaminoglycan (s-GAG) molecules in differentiated and non-differentiated dECM (Figure 3D). Furthermore, s-GAG quantification was performed between the two samples (Figure 4**-3 E**). The staining demonstrated a robust and intense blue colour, indicating the accumulation of s-GAG molecules in the differentiated sample. The findings from s-GAG quantification along with Alcian blue staining results suggest successful chondrogenic differentiation of the ATDC5 cells, as s-GAGs are a hallmark of cartilage tissue and play a crucial role in maintaining its structure and function. The ability of the ATDC5 cells to differentiate into chondrocyte-like cells and produce s-GAGs has been well-documented in various studies[27,28,44]. Moreover, the significant increase in s-GAG synthesis is in line with previous research that highlights the importance of insulin, transferrin and selenium in driving chondrogenesis[28]. The observed increase in s-GAG synthesis can be attributed to the activation of various intracellular signalling pathways and transcription factors involved in chondrogenesis. These include *SOX9*, a master regulator of chondrogenic differentiation, which plays a crucial role in the synthesis of cartilage-specific extracellular matrix components, including s-GAGs[45]. Differentiation likely upregulated the expression and activity of *SOX9*, leading to enhanced s-GAG synthesis in the ATDC5 cells.

**Figure 3.**
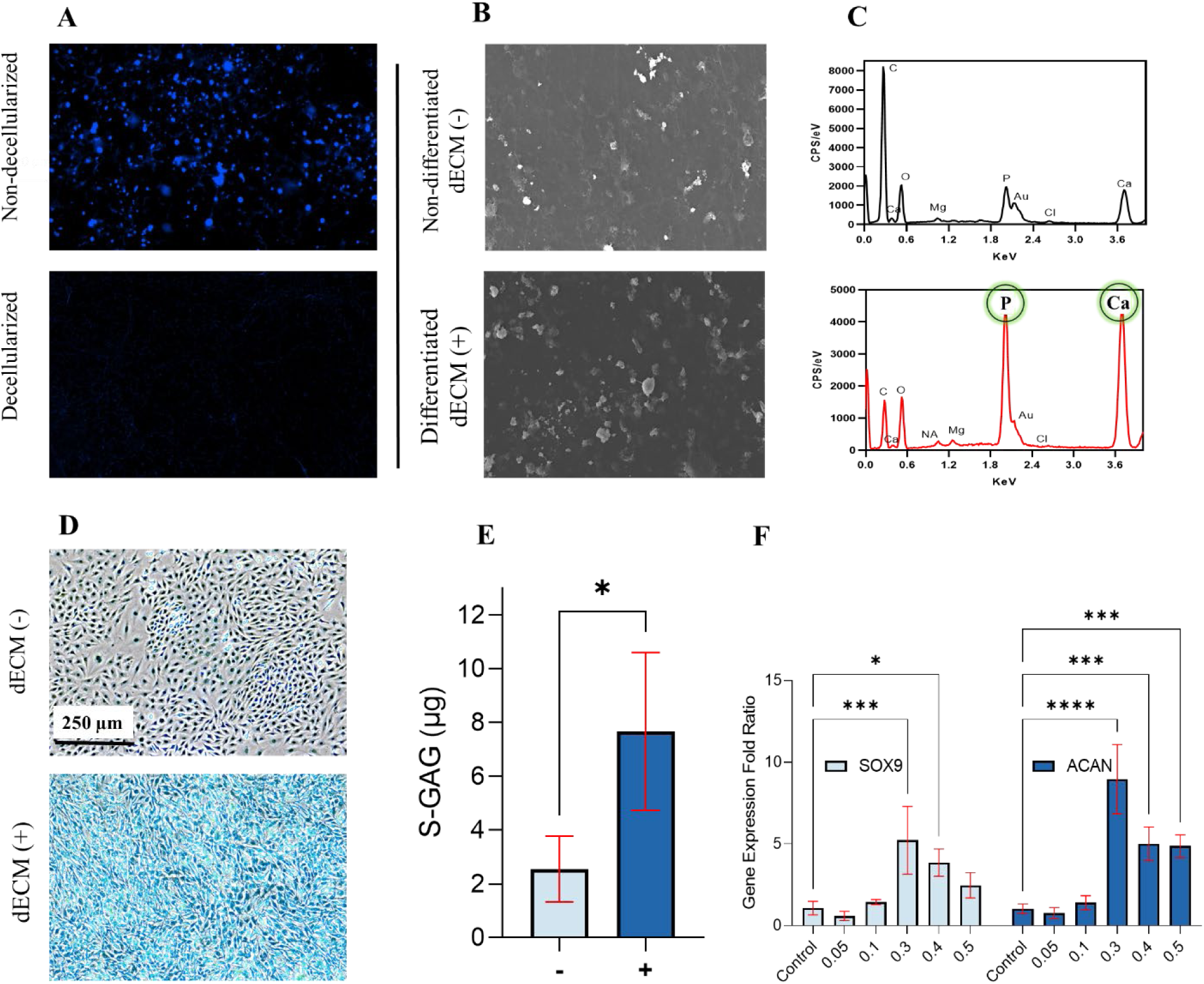
Characterization of decellularized extracellular matrix (dECM) obtained from differentiated pre-chondrogenic cells (ATDC5). A) DAPI staining of ATDC5 cells before (up) and after (down) decellularization, indicating successful removal of cellular material. B) SEM images comparing dECM derived from differentiated (+) and non-differentiated (-) ATDC5 cells, showing structural differences. C) EDX analysis of the two groups, revealing relatively higher amount of P and Ca^+2^ in the differentiated dECM samples. D) Alcian Blue staining of the ATDC5 cells, illustrating presence of s-GAG molecules in the differentiated dECM. E) Comparison of differentiated and non-differentiated ATDC5 in s-GAG synthesis (n=3). ATDC5 showed significantly higher synthesis of s-GAGs upon introduction to chondrogenic differentiating media. F) RT-qPCR analysis of chondrogenic markers (*SOX9* and *ACAN*) in hADSCs treated with varying concentrations of CdECM (w/v), normalized to the *GAPDH* reference gene (n=3). 0.3% concentration showed the highest gene expression. (* p<0.05, *** p<0.001, **** p<0.0001).

**Figure 4.**
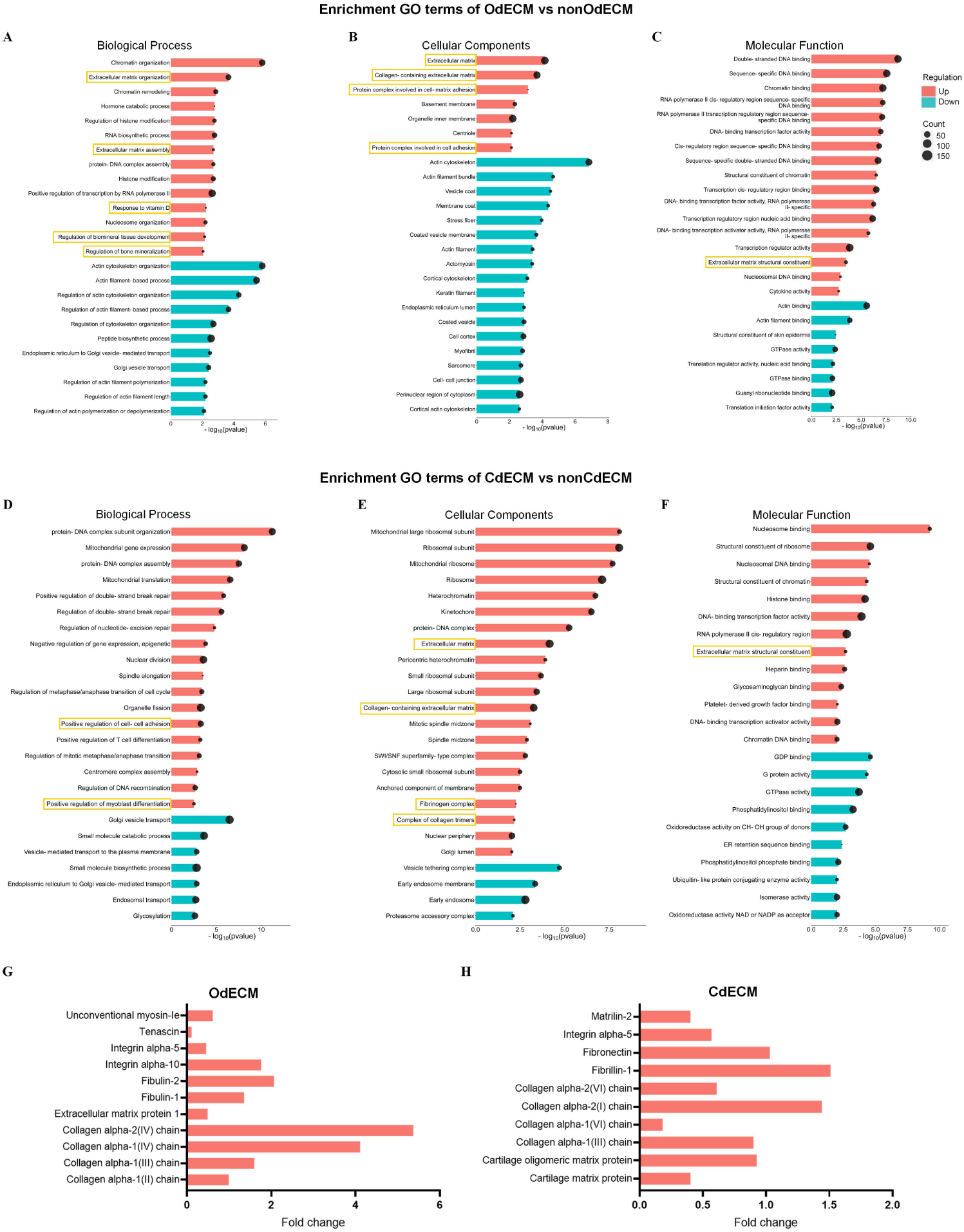
Proteomic Analysis of OdECM and CdECM. A-C) Functional enrichment analysis of OdECM compared to normal dECM including distinct upregulated and downregulated proteins in biological processes (A), cellular components (B) and molecular functions (C). D-F) Functional enrichment analysis of Chondrogenic ECM compared to normal ECM including distinct upregulated and downregulated proteins in biological processes (D), cellular components (E) and molecular functions (F). G) Change of upregulated bone ECM proteins in Osteogenic ECM compared to normal ECM. H) Change of upregulated bone ECM proteins in Chondrogenic ECM compared to normal ECM. GO terms with top 25 enriched genes was shown in each functional enrichment graph. Gene ontology terms related to bone or cartilage development and ECM functions are marked with yellow boxes.

RT-qPCR analysis was also conducted to investigate the expression of chondrogenic markers, specifically *SOX9* and *ACAN*, in hADSCs treated with varying concentrations of CdECM derived from ATDC5 cells (Figure 3F). *SOX9* and *ACAN* are two critical genes involved in chondrogenesis, the process of cartilage formation and development[45]. *SOX9* encodes a transcription factor that plays a central role in regulating the differentiation and maturation of chondrocytes. It is essential for the expression of numerous genes involved in cartilage development, including *ACAN*[45]. *ACAN* encodes a large proteoglycan that constitutes a major component of ECM in cartilage tissue[46]. It provides structural integrity to cartilage and is crucial for maintaining its compressive strength and hydration properties. The pathways involved in the regulation of *SOX9* and *ACAN* during chondrogenesis are complex and tightly coordinated. Several signalling pathways and transcription factors contribute to their expression and activity. One of the key pathways involved is the Wnt/β-catenin signalling pathway[47]. During the early stages of chondrogenesis, Wnt signalling is suppressed, allowing *SOX9* expression and subsequent chondrocyte differentiation[47]. Later, as chondrocytes mature, Wnt signalling is activated, leading to the suppression of *SOX9* and the promotion of hypertrophic differentiation and enlargement.

The RT-qPCR analysis revealed that a 0.3% concentration of CdECM resulted in the highest expression of chondrogenic markers in hADSCs. This concentration was used for all future experiments. This finding indicates that the dECM derived from differentiated ATDC5 cells contains chondrogenic factors that enhance the chondrogenic potential of hADSCs and influence the behaviour of stem cells towards a chondrogenic phenotype.

### 3.4 Proteomics analysis further establishes the upregulation of osteogenic and chondrogenic proteins associated with OdECM and CdECM, respectively

To investigate the change of OdECM and CdECM proteins compared to ECM produced by undifferentiated precursor cells, a quantitative proteomic analysis was performed to identify the protein profiles of OdECM/nonOdECM and CdECM/nonCdECM. For the bone ECM, 2447 proteins were identified in both OdECM and nonOdECM samples, with 15 unique proteins in OdECM and 61 unique proteins in nonOdECM respectively. Functional enrichment analysis of all proteins present in both OdECM and nonOdECM showed upregulation of bone ECM-related gene ontology (GO) terms including ECM production, cell adhesion and bone mineralization (Figure 4A-C). These enriched GO terms demonstrated the increased osteoinductive potential of ECM after osteogenic differentiation in MC3T3-E1 pre-osteoblast cells. Similarly, the proteins identified in CdECM also showed an enhanced chrondroinductive extracellular environment compared to nonCdECM. The CdECM proteins were enriched in positive regulation of ECM formation including cell to cell adhesion, collagen-containing ECM components and fibrinogen production (Figure 4D-F). To further confirm the osteogenic and chondrogenic profiles, change in typical bone and cartilage ECM structural proteins was analyzed using the quantitative data from ECM proteomics. Figure 4G showed more than 2-fold increase of Type I collagen and a minor increase of fibulin and integrin alpha 10 in OdECM [48]. In Figure 4H, vital cartilage ECM structural proteins including fibronectin and cartilage matrix protein were also increased[49]. In conclusion, enhanced osteoinductive and chrondroinductive potential was identified in OdECM and CdECM respectively by proteomic analysis.

### 3.5 OdECM and CdECM can be integrated in GelMA pregel and 3D printed using a light-based bioprinter

Hydrogels containing OdECM (GelO) and CdECM (GelC) were printed using a Digital Light Processing (DLP)-based printer. DLP printing utilizes light (often blue or UV light) to fabricate hydrogels with high precision and resolution. This technique works by projecting light patterns onto a liquid bioink, solidifying it layer by layer to form the desired 3D structure[50]. The ability to create patient-specific geometries is a significant advantage of 3D printing in tissue engineering. This customization allows for the fabrication of hydrogels that precisely match the defect or injury site, optimizing tissue regeneration outcomes[51]. Figure 5A shows the printer and the computer aided design used for fabricating the gels that were used for the mechanical characterization of the hydrogels. The corresponding hydrogels shown in the figure, demonstrates the capability of the printing technique to produce precise and reproducible geometries, which is critical for fabricating tissue engineering scaffolds. Scanning Electron Microscopy (SEM) images were obtained to investigate the microstructure of the hydrogels (Figure 5B). The SEM analysis revealed macroporous void spaces in all the hydrogel formulations, indicative of their three-dimensional network structure. The interconnected porous architecture of the hydrogels provides a favourable environment for cell infiltration, proliferation, and tissue regeneration. A reduction in porosity was evident upon the incorporation of dECM into the hydrogels. This finding suggests that the incorporation of dECM leads to a more densely crosslinked network, potentially enhancing the overall mechanical properties of the hydrogels. The increased crosslinking density with dECM might be attributed to enhanced interaction between dECM particles and the polymer network of GelMA due to the presence of additional functional groups and reactive sites. Additionally, it is plausible that the introduction of dECM reduced the penetrance of blue light during the photo-crosslinking process, resulting in a tighter and less porous network formation.

**Figure 5.**
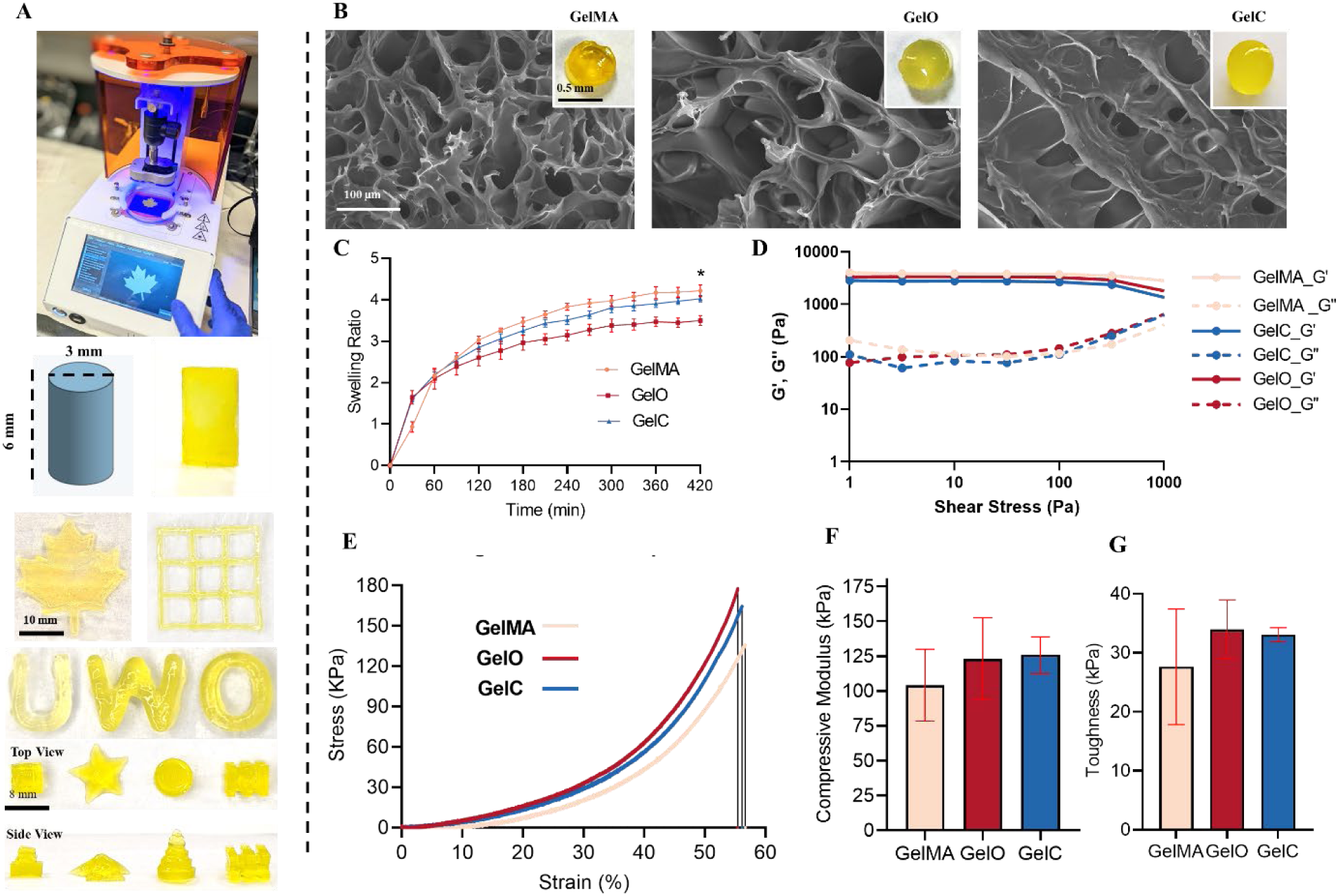
Physicochemical And Mechanical Characterization of GelO and GelC. A) Schematic of DLP printer Lumen X programed with a cylindrical Onsahpe design and its corresponding GelC hydrogel for mechanical testing (diameter: 3mm, height: 6mm). Other hydrogels have been 3D-printed to showcase sensitivity of the printing technique. B) SEM images have been displayed revealing macroporous void spaces in the hydrogel. SEM visually showed the reduction in porosity upon the introduction of dECM. C) Graph exhibits the swelling profile of GelMA, GelO and GelC (n=5). Swelling of GelO and GelC was reduced significantly compared to GelMA. D) Rheology experiment depicting the storage modulus (G’) and loss modulus (G“) profiles of GelMA, GelO, and GelC gels. Replicates (n=3) of 6 cm x 6 cm x 6 cm cylindrical gels were printed and hydrated in PBS for 5 minutes (only one replicate is shown). stress sweep was performed at 37 °C from 0.1 to 104 Pa. The results demonstrate a very similar profile among the groups, with GelMA exhibiting a slightly higher storage modulus compared to GelO and GelC. E) A representative stress vs strain plot of formulated hydrogels obtained under uniaxial compression tests up to 0.6 mm/mm strain. F) Comparison between the compressive moduli of GelMA, GelO and GelC hydrogels. An increase in compressive strength was observed with the incorporation of dECM. Compressive modulus was determined from the slope of the stress-strain plot in the region of 1-15% strain. Results are shown as mean ± standard deviation (n = 5). G) Toughness test results showing a comparison of GelMA, GelO, and GelC gels. Toughness values (kPa) were measured to assess the resistance to fracture upon compressive force. The data reveals that the inclusion of dECM slightly increased the toughness in comparison to the samples without dECM (n=3). (* p<0.05)

### 3.6 Incorporation of OdECM and CdECM reduces swelling and degradation of the hydrogels

The swelling profiles of hydrogels were investigated to understand their water uptake capacities (Figure 5C). The ability of hydrogels to absorb and retain water is essential for maintaining a hydrated environment within the scaffold, supporting cell growth, and facilitating nutrient diffusion. The results demonstrate that GelO and GelC exhibited significantly reduced swelling compared to GelMA. The reduction in swelling indicates a tighter network structure in GelO and GelC, corroborating the SEM observations. The diminished swelling behaviour may be attributed to the presence of dECM, which could restrict water penetration and result in a more compact hydrogel structure. Additionally, the dECM molecules may sterically hinder the access of water molecules to the hydrogel matrix, leading to reduced swelling. We also investigated the degradation behaviour of the hydrogels on day 10. Degradation studies provide valuable insights into the hydrogel’s ability to maintain structural integrity and its susceptibility to breakdown over time. The percentage of degradation was measured as 2.012 ± 0.38 for GelMA, 1.69 ± 0.10 for GelO, and 1.87 ± 0.38 for GelC. The results indicated that GelO and GelC hydrogels exhibited slightly lower degradation rates compared to GelMA. This reduction in degradation may be attributed to the incorporation of dECM, which could act as a protective barrier. Similar to swelling results, the presence of dECM particles may also create additional crosslinks, further stabilizing the hydrogel structure and contributing to its enhanced resistance against degradation.

### 3.7 OdECM and CdECM do not alter the rheological properties of the hydrogels

Rheological experiments were conducted to assess the viscoelastic properties of GelMA, GelO, and GelC hydrogels (Figure 5D). The storage modulus (G’) and loss modulus (G“) profiles were measured by subjecting the cylindrical gels to a stress sweep at 37 °C. The rheological behaviour of hydrogels is crucial as it reflects their mechanical stability, ability to withstand external forces, and their potential as tissue engineering scaffolds. Remarkably, the three hydrogel formulations exhibited very similar rheological profiles. GelMA displayed a slightly higher storage modulus compared to GelO and GelC, suggesting that it possesses greater elastic behaviour and resistance to deformation. The slight variation in viscoelastic behaviour may arise from differences in the molecular weight and density of the hydrogel networks.

### 3.8 OdECM and CdECM enhances the mechanical strength of the hydrogels

Uniaxial compression tests were conducted to analyze the compressive strength of the formulated hydrogels (Figure 5E). The stress vs. strain plots demonstrated the mechanical response of the hydrogels under compressive force up to 0.6 mm/mm strain. The compressive strength of hydrogels is essential in supporting and maintaining tissue structure under mechanical loading. Figure 5F presents a comparison of the compressive moduli of GelMA, GelO, and GelC hydrogels within the linear section of the stress-strain curve. The data revealed an increase in compressive strength with the incorporation of dECM. GelO and GelC hydrogels, with dECM, exhibited enhanced mechanical stiffness compared to GelMA, suggesting that the presence of dECM contributes to the overall reinforcement of the hydrogel matrix. The integration of dECM potentially creates a composite-like structure, combining the mechanical properties of both components and resulting in a stiffer and more robust hydrogel.

Next, toughness was determined to assess the ability of the hydrogels to resist fracture upon being subjected to compressive force (Figure 5G). The results showed that the inclusion of both OdECM and CdECM slightly increased the toughness of GelO and GelC hydrogels compared to their counterparts without dECM. Toughness is a critical mechanical property in tissue engineering as it reflects the energy required to induce failure in the hydrogel. The improved toughness can be attributed to structural integrity conferred by the presence of dECM, which enhances the hydrogel’s resistance to fracture and deformation. The improved toughness indicates that the hydrogels with dECM are better able to withstand mechanical challenges *in vivo*, making them more suitable candidates for non-load-bearing applications.

The observed improvements in mechanical properties, coupled with the ability to control structural characteristics through 3D printing, underscore the potential of GelO and GelC hydrogels as promising biomaterials for osteochondral repair applications. Continued investigation into the effects of dECM concentration and other modifications on hydrogel properties will help refine and tailor these hydrogels for specific bone and cartilage tissue engineering applications.

### 3.9 Cytocompatible GelO is capable of inducing osteogenic differentiation in hADSCs

After establishing a formulation for Gel, we aimed to evaluate the cytocompatibility of the bioactive hydrogel by investigating the organization and morphology of cytoskeleton and actin filaments, as well as cell viability and metabolic activity (Figure 6A). Fluorescent staining using Phalloidin (green) and DAPI (blue) was employed to assess the structural integrity of hADSCs cultured on thin layers of GelMA and GelO for 24 hours.

**Figure 6.**
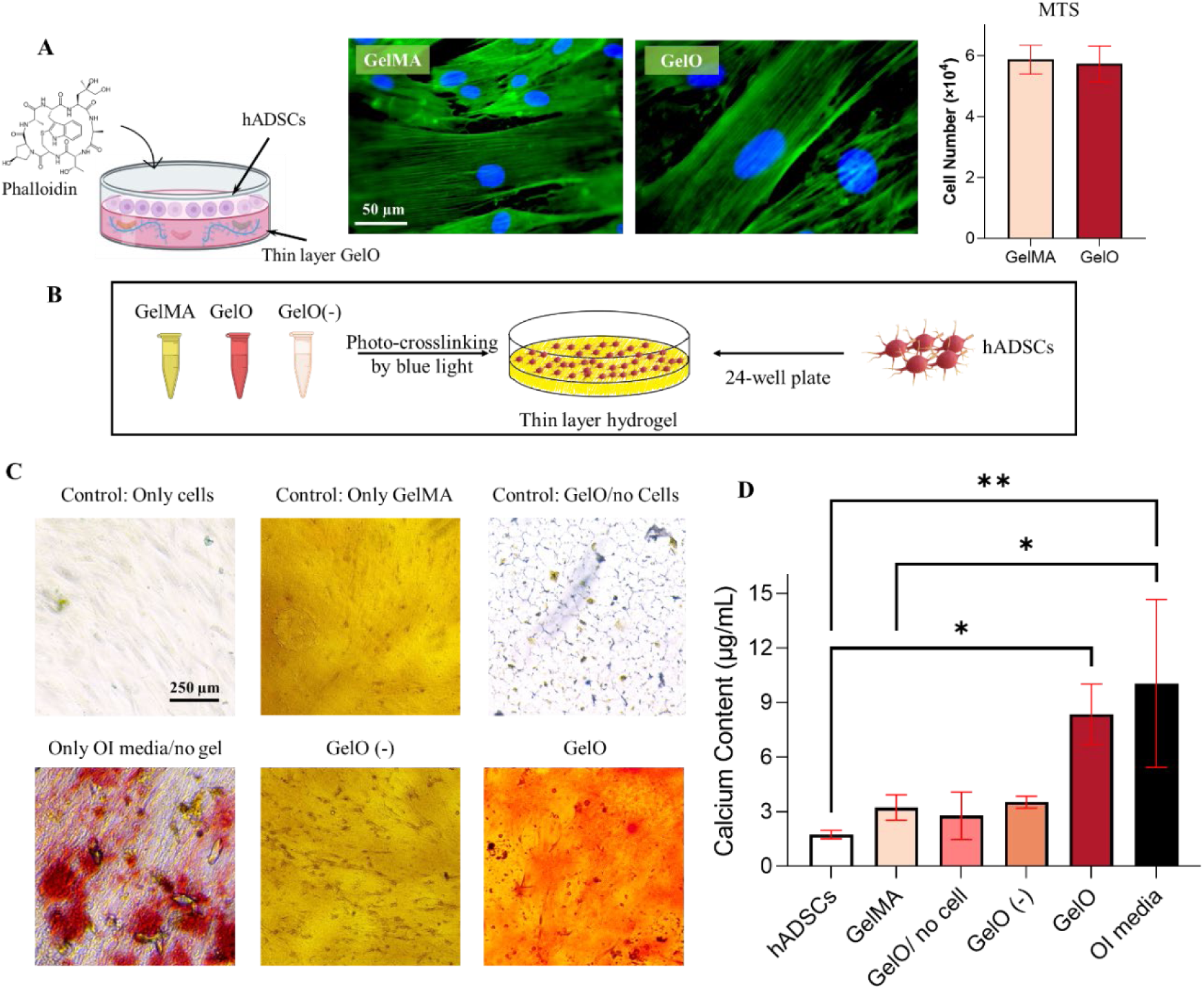
Comparative Analysis of Stem Cell Growth and Osteogenic Differentiation on GelMA and GelO. A) **Phalloidin/DAPI Staining and MTS Assay Results**: Fluorescent staining using Phalloidin (green) and DAPI (blue) was performed on hADSCs cultured on thin layer GelMA and GelO for 24 hours. The staining revealed that the organization and morphology of the cytoskeleton and actin filament (green) and cell nuclei (blue) stayed intact. The corresponding MTS assay was conducted to assess the metabolic activity and cell viability of the stem cells. The bar graph represents the cell number, demonstrating the relative cell viability between the two groups. B) **Alizarin Red Staining Experiment on hADSCs**: An illustration depicts the experimental setup for the Alizarin Red staining assay performed on hADSCs. The cells were cultured on a thin layer of hydrogels for 21 days to induce osteogenic differentiation. C) **Microscopy Images of Alizarin Red Stained hADSCs**: Microscopy images exhibit the results of Alizarin Red staining on hADSCs. The sample treated with osteoinductive media (OI), containing dexamethasone in DMEM, acted as positive group while all the other groups were cultured in ADSC-BM media. GelO samples showed visible mineralization that indicates successful osteogenic differentiation. The GelO (-) group refers to the hydrogel that contains 0.4% (w/v) dECM derived from non-differentiated MC3T3-E1-E1 cells. D) **Quantification of Alizarin Red Staining**: Quantitative analysis was performed to measure the dye extraction from the Alizarin Red staining assay. The results demonstrated that hADSCs treated with GelO exhibited significantly higher levels of staining compared to controls, suggesting enhanced osteogenic differentiation.

The staining results revealed that GelMA and GelO hydrogels supported the preservation of the organization and morphology of the cytoskeleton and actin filaments, as indicated by the intense green fluorescence. This suggests that the hydrogels did not disrupt the structural integrity of hADSCs, demonstrating their biocompatibility. Additionally, the blue fluorescence from DAPI staining indicated that the cell nuclei remained intact, further confirming the absence of genotoxic effects induced by the hydrogels. The maintenance of cellular architecture is of utmost importance, as it directly influences cell behaviour, including adhesion, migration, and differentiation. To assess the metabolic activity and cell viability of hADSCs cultured on GelO, an MTS assay was conducted, as displayed in the bar graph (Figure 6A). The data indicated that both GelMA and GelO hydrogels were able to support the survival and proliferation of hADSCs, as evidenced by the similar metabolic activity observed in both groups. GelMA hydrogels have been recognized for their potential in wound healing, drug delivery, biosensing, and tissue regeneration due to their biocompatibility and tunable physical properties[52]. Our findings showed the addition of dECM into GelMA did not affect the cytocompatibility of hydrogels. This suggests that these hydrogels can serve as suitable substrates for supporting osteogenic and chondrogenic differentiation.

After demonstrating cytocompatibility, we evaluated the osteogenic potential of the developed GelO. Figure 6B illustrates the corresponding experimental setup, where Alizarin red stain was used to determine the deposition of mineral. The hADSCs were cultured on a thin layer of hydrogels for 21 days to induce osteogenic differentiation. Micrographs of Alizarin Red stain has been displayed in Figure 6C. Here, the positive group designated as OI was treated with osteoinductive media containing dexamethasone in DMEM. OI served as a positive control that induce osteogenic differentiation with the help of drugs. We chose to use DMEM as basal media for positive control because it has been reported that stem cells show a maximal differentiation in DMEM supplemented with appropriate drugs or serums compared to other types of media[26,53–55]. Intense red and blue colours in OI were observed due to the induction of differentiation by dexamethasone. The other groups were cultured in hADSC basal media without the osteoinductive factors. It can be observed that the hADSCs grown on GelO exhibited visible mineralization, indicating successful osteogenic differentiation (Figure 6C). On the other hand, the GelO (-) group, which refers to the hydrogel containing 0.4% (w/v) dECM derived from non-differentiated MC3T3-E1 cells, showed similar colour intensity to the GelMA control group. This finding suggests that differentiation of pre-osteoblastic cells is essential to harvest osteoinductive dECM. The quantitative analysis of the Alizarin Red staining results is depicted in Figure 6D. The staining levels were measured by extracting the dye and quantifying its absorbance at 405 nm. The results demonstrated that hADSCs treated with GelO hydrogel exhibited significantly higher levels of staining compared to the control groups, indicating enhanced osteogenic differentiation. The GelO hydrogel likely provides a conducive microenvironment for hADSCs, facilitating their commitment toward the osteogenic lineage.

### 3.10 Cytocompatible GelC is capable of inducing chondrogenic differentiation in hADSCs

Figure 7A demonstrates the cytocompatibility of GelC using hADSCs. Phalloidin and DAPI stains demonstrate that GelC does not disrupt the structural integrity of hADSCs grown on the hydrogels. Further, MTS assay indicates that GelC supports cell survival and growth. Next, the chondrogenic potential of GelC was determined. Here too, the cells were cultured on a layer of GelC for 21 days. The cells were then stained with Alcian blue, as shown in the illustration for experimental setup (Figure 7B). A chondroinductive media containing DMEM/F12 was used as a positive control (CI). Intense blue color was observed in the positive control because of the inevitable differentiation from the use of commercial chondroinductive drugs. the hADSCs cultured on GelC hydrogel showed a prominent and intense blue coloration, indicative of a higher synthesis of sulfated glycosaminoglycans (s-GAGs). This robust s-GAG synthesis observed in the GelC group provides strong evidence of successful chondrogenic differentiation, implying the formation of cartilage-like structures (Figure 7C). In contrast, the GelC (-) group, which consisted of the hydrogel containing 0.3% (w/v) dECM derived from non-differentiated ATDC5 cells, displayed a colour intensity similar to that of the GelMA control group. This finding suggests that the differentiation of pre-chondrogenic cells is crucial for obtaining chondroinductive dECM, as the GelC (-) group lacks the distinct blue coloration associated with increased s-GAG synthesis., Figure 7D presents the quantitative analysis derived from the Sulfated-Glycosaminoglycans Assay Kit. This assay efficiently detects all s-GAGs, including heparan sulfate, chondroitin sulfate, and keratan sulfate. The principle of the assay involves the interaction of a cationic dye, 1,9-dimethylmethlene Blue (DMMB), with the highly negatively charged s-GAGs, resulting in the formation of a coloured product. The absorbance of this coloured signal is measured at 525 nm and is directly proportional to the concentration of sulfated GAGs present in the sample. The outcome of the quantitative analysis clearly indicated that hADSCs treated with the GelC hydrogel exhibited significantly higher levels of staining compared to the control groups. This observation strongly suggests enhanced chondrogenic differentiation in the GelC group. The GelC hydrogel likely provides a favourable microenvironment that promotes hADSCs’ commitment toward the chondrogenic lineage, leading to the augmented synthesis of sulfated GAGs.

**Figure 7.**
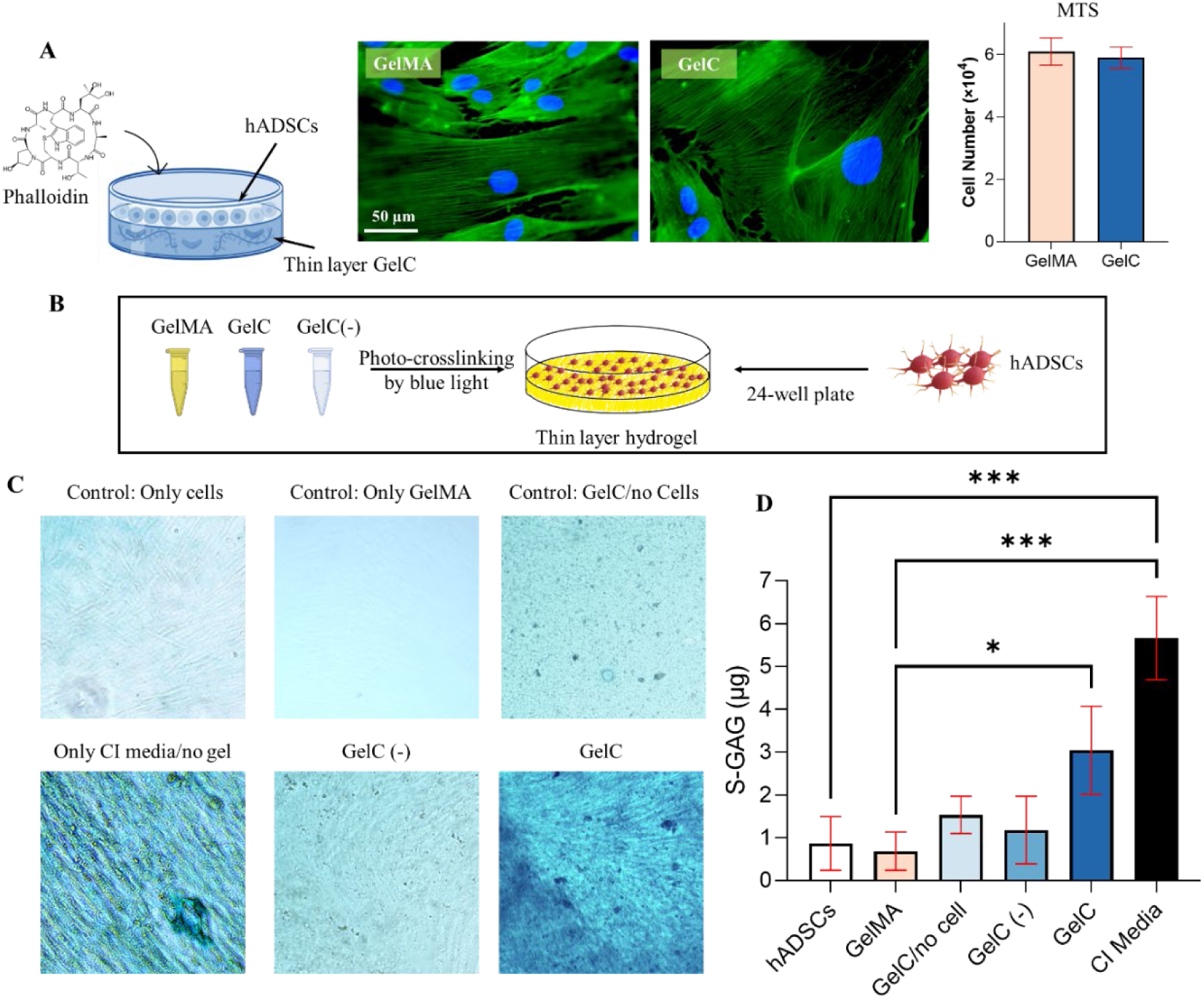
Comparative Analysis of Stem Cell Growth and Chondrogenic Differentiation on GelMA and GelC. A) **Phalloidin/DAPI Staining and MTS Assay Results**: Phalloidin/DAPI staining was performed on stem cells cultured on GelMA and GelC after 24 hours to visualize the preservation of cytoskeleton (green) and cell nuclei (blue). The corresponding MTS assay was conducted to assess the metabolic activity and cell viability of the stem cells. The bar graph presents the cell number values obtained, indicating similar cell viability between the two groups. B) **Illustration of Alcian Blue Staining Experiment on hADSCs**: An illustration depicts the experimental setup for the Alcian Blue staining experiment conducted on hADSCs grown on a thin layer of GelC for 21 days to induce chondrogenic differentiation. **C) Microscopy Images of Alcian Blue Stained hADSCs**: Microscopy images display the results of Alcian Blue staining on hADSCs. The samples treated with chondroinductive media (CI), containing dexamethasone and ITS serum in DMEM/F12, acted as positive group while all the other groups were cultured in ADSC-BM media. GelC samples exhibited positive staining for sulfated glycosaminoglycans (s-GAG), indicating successful chondrogenic differentiation. GelC (-) represents the that contains 0.3% (w/v) dECM derived from non-differentiated ATDC5 cells. D) **Quantification of s-GAG**: Quantitative analysis was conducted to measure the sulfated glycosaminoglycan (s-GAG) content with the use of Sulfated-Glycosaminoglycans Assay Kit. hADSCs treated with GelC hydrogel demonstrated significantly higher levels of s-GAG compared to the control groups, suggesting enhanced chondrogenic differentiation.

### 3.11 GelO and GelC together are capable of inducing osteochondrogenesis in hADSCs

Osteogenesis and chondrogenesis are vital processes in osteochondral tissue repair. The ability to promote the differentiation of stem cells into osteoblasts and chondrocytes using hydrogel-based treatments is of great interest and is the final goal of this study. Here, we sought to evaluate the effectiveness of three different hydrogel combinations (Gel-GelO, Gel-GelC, GelO-GelC) in inducing osteogenic and chondrogenic differentiation (Figure 8). The experimental design involved a side-by-side formation of thin-layer hydrogel treatments, and gene expression analysis was used to quantify the differentiation outcomes.

**Figure 8.**
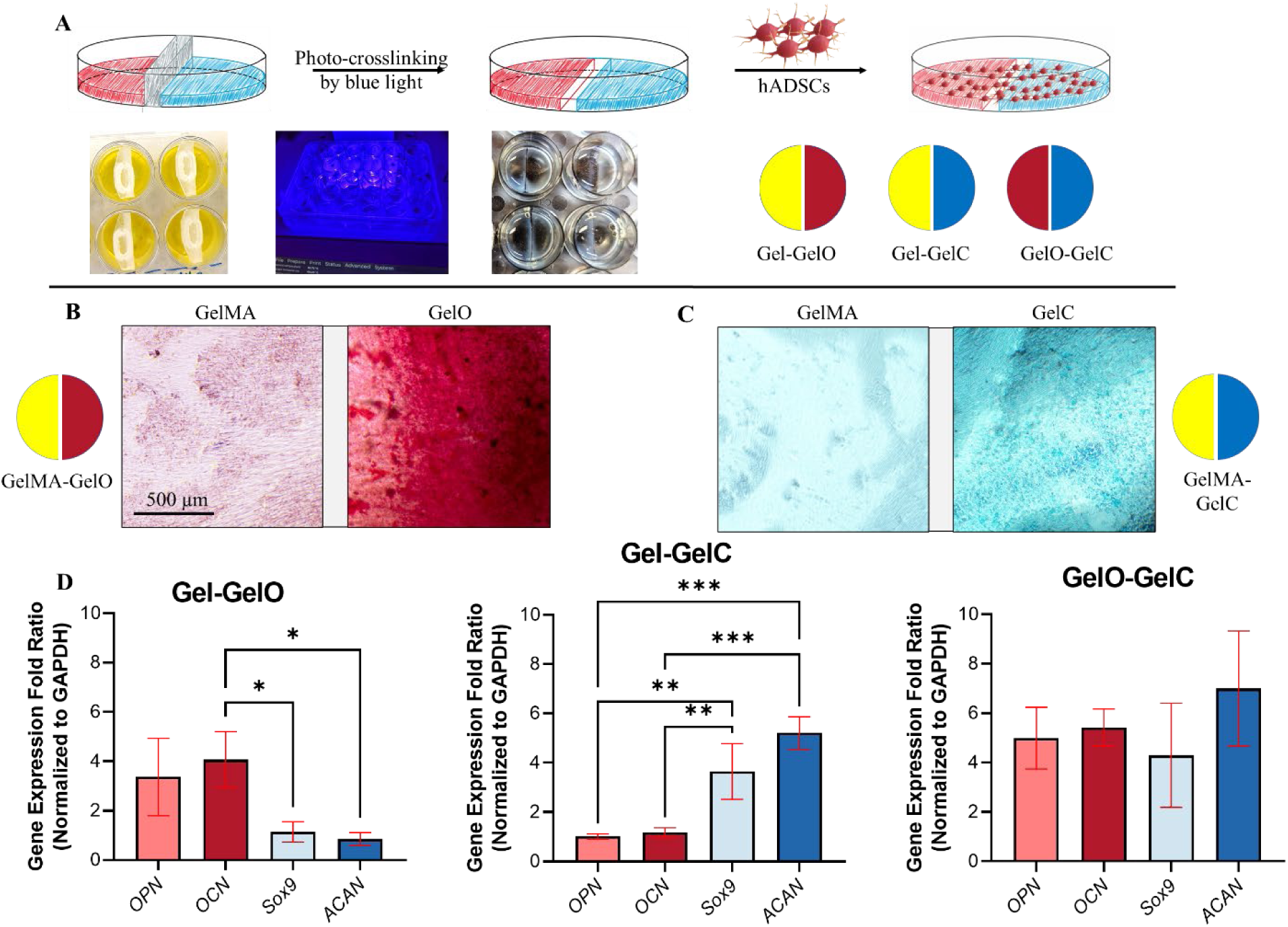
Comparative Analysis of Osteogenesis and Chondrogenesis on Combination of Thin Layer Hydrogel Treatments. A) The cartoon depicts the experimental design for side-by-side thin layer hydrogel treatments in a 24-well plate. A plastic tight wall was inserted in the middle of the wells, creating separate compartments for GelMA, GelO, and GelC. Blue light was applied for photo-crosslinking of the hydrogel premixes. hADSCs were seeded on top of the hydrogels and cultured for 21 days to analyze gene expression. B) Alizarin Red staining was performed on Gel-GelO samples to assess osteogenic differentiation. The bright red color observed in the GelO side indicated the presence of calcium deposits, indicative of successful osteogenesis. In contrast, the GelMA side exhibited limited red staining, suggesting minimal osteogenic differentiation. C) Alcian Blue staining was conducted on Gel-GelC samples to evaluate chondrogenic differentiation. The presence of bright blue color in the GelC side indicated the deposition of sulfated proteoglycans, a characteristic feature of chondrogenesis. Conversely, the GelMA side showed minimal blue staining, indicating limited chondrogenic differentiation. D) RT-qPCR analysis was performed to quantify the expression levels of osteogenic markers (OPN and OCN) and chondrogenic markers (SOX9 and ACAN) in hADSCs treated with Gel-GelO, Gel-GelC, and GelO-GelC. The gene expression levels were normalized to the reference gene GAPDH. The Gel-GelO group exhibited significantly higher expression levels of osteogenic markers, indicating enhanced osteogenic differentiation compared to the other groups. Similarly, the Gel-GelC group demonstrated higher expression levels of chondrogenic markers, suggesting enhanced chondrogenic differentiation. Notably, the GelO-GelC group exhibited a remarkable 4 to 6-fold upregulation of both osteogenic and chondrogenic markers, indicating synergistic effects of GelO and GelC in promoting both lineages (* p<0.05, ** p<0.01, *** p<0.001).

The setup included a 24-well plate with separate compartments created by a plastic tight wall inserted in the middle (as shown in Figure 8A). The three hydrogel types used were GelMA, GelO, and GelC. Blue light was applied to photo-crosslink the hydrogel premixes. hADSCs were seeded on top of the hydrogels and cultured for 21 days to allow for osteogenic and chondrogenic differentiation. For the assessment of osteogenesis, Alizarin Red staining was performed on Gel-GelO samples (Figure 8B). The GelO side displayed a bright red colour, indicating the presence of calcium deposits. In contrast, the GelMA side exhibited limited red staining, suggesting minimal osteogenic differentiation. This result indicates that GelO was more effective than GelMA in promoting the osteogenic differentiation of hADSCs. To evaluate chondrogenesis, Alcian Blue staining was conducted on Gel-GelC samples (Figure 8C). The GelC side displayed a bright blue colour, indicating the deposition of sulfated proteoglycans, a characteristic feature of chondrogenesis. On the other hand, the GelMA side exhibited limited blue staining, suggesting limited chondrogenic differentiation. This finding suggests that GelC was more effective in promoting chondrogenic differentiation compared to GelMA within the same well.

To quantify the expression levels of osteogenic and chondrogenic markers, RT-qPCR analysis was performed on hADSCs treated with different hydrogel combinations (Figure 8D). The gene expression levels were normalized to the reference gene *GAPDH*. The Gel-GelO group showed significantly higher expression levels of osteogenic markers, *OPN* and *OCN*, indicating enhanced osteogenic differentiation compared to the other groups. Furthermore, the Gel-GelC group demonstrated higher expression levels of chondrogenic markers, *SOX9* and *ACAN*, suggesting enhanced chondrogenic differentiation compared to the other groups. These results are consistent with the findings from Alizarin Red and Alcian Blue staining, indicating that GelO and GelC effectively promote lineage-specific differentiation in hADSCs.

The most intriguing finding of this study was observed in the GelO-GelC group. This group exhibited a remarkable 4 to 6-fold upregulation of both osteogenic and chondrogenic markers, indicating synergistic effects of GelO and GelC in promoting both lineages. The increased expression of *OPN* and *OCN* suggests that the combination of GelO and GelC not only enhances osteogenesis but also induces an osteoinductive microenvironment that promotes the commitment of hADSCs towards osteoblast-like cells. Similarly, the upregulation of chondrogenic markers, indicates that the combination of GelO and GelC synergistically induces chondrogenic differentiation in hADSCs. The observed synergistic effect of GelO and GelC can be attributed to their distinct mechanisms of action. During early osteogenesis, cells are known to release osteoinductive factors such as Runx2 that stimulate the differentiation of hADSCs into osteoblasts[56]. Runx2 is one of the earliest transcription factors upregulated during osteoblast differentiation, and its expression is essential for the commitment of MSCs to the osteoblastic lineage. Runx2 acts as a transcriptional activator, binding to specific DNA sequences in the regulatory regions of numerous osteogenic genes including *OCN* and *OPN* to stimulate their expression[56]. As osteoblasts mature Runx2 levels decrease, and this fluctuation also affects chondrogenesis. In the early stages of chondrogenic differentiation, *Runx2* expression is upregulated in stem cells as they condense to form chondroprogenitor cells[57]. *Runx2* is also involved in the transition of chondrocytes to hypertrophic chondrocytes, a process that occurs during endochondral ossification[57]. In this process, chondrocytes in the center of the cartilage model enlarge and undergo mineralization, leading to the formation of the primary ossification center. *Runx2* expression is upregulated in hypertrophic chondrocytes and plays a role in the mineralization of the cartilage matrix[57]. However, as chondrogenesis progresses, *Runx2* expression is downregulated in committed chondrocytes. Therefore, the combination of GelO and GelC hydrogels likely creates a multifunctional microenvironment that enhances the expression of lineage-specific transcription factors, promoting both osteogenesis and chondrogenesis. The observed synergistic effect of GelO and GelC in promoting both osteogenesis and chondrogenesis aligns with the notion that combining multiple cues can lead to enhanced differentiation outcomes. Previous studies have shown that biomaterials can act synergistically to regulate stem cell fate. For example, researchers fabricated a biphasic hydrogel called CAN-PAC to facilitate the regeneration of osteochondral defects[58]. In an experimental rabbit model, bilayer hydrogels were applied to the defect sites. The regenerated tissues exhibited newly formed transparent cartilage and repaired subchondral bone, providing evidence of the hydrogel’s efficacy in promoting osteochondral defect repair[58].

This comparative analysis of thin-layer hydrogel treatments demonstrated the differential abilities of GelO and GelC in promoting osteogenesis and chondrogenesis. The GelO-GelC combination displayed a remarkable synergistic effect in promoting both lineages, making it a promising candidate for future osteochondral repair applications. Further investigations are warranted to understand the underlying mechanisms responsible for this synergistic effect and to optimize the hydrogel composition and culture conditions.

### 3.12 Covalently adhering GelO and GelC to form bioactive plug for repairing small osteochondral defects

The process of conjugating GelMA hydrogels through EDC/NHS coupling for osteochondral plug design offers a promising approach to osteochondral tissue repair. In Figure 9A, a detailed illustration showcases the step-by-step process of conjugating two GelMA hydrogels through EDC/NHS coupling. EDC acts as a crosslinker that activates carboxylic acid groups on the GelMA hydrogels, generating highly reactive O-acylisourea intermediates[59]. NHS, on the other hand, functions as a stabilizing agent for the O-acylisourea intermediates. The O-acylisourea intermediate readily reacts with primary amines present on the second GelMA chains, resulting in amide bond formation, and effectively conjugates the GelMA hydrogels[59]. The use of EDC/NHS coupling provides several advantages in the construction of the multilayer hydrogel. First, the process ensures a strong and stable covalent linkage between the hydrogels, enhancing the structural integrity of the composite material[60]. This is critical for its successful implantation and long-term performance in the challenging osteochondral defect environment. Second, the covalent bonds formed via EDC/NHS coupling do not adversely affect the biocompatibility of the hydrogel as studies have used this technique in *in vivo* and *in vitro* experiments[61–63].

**Figure 9.**
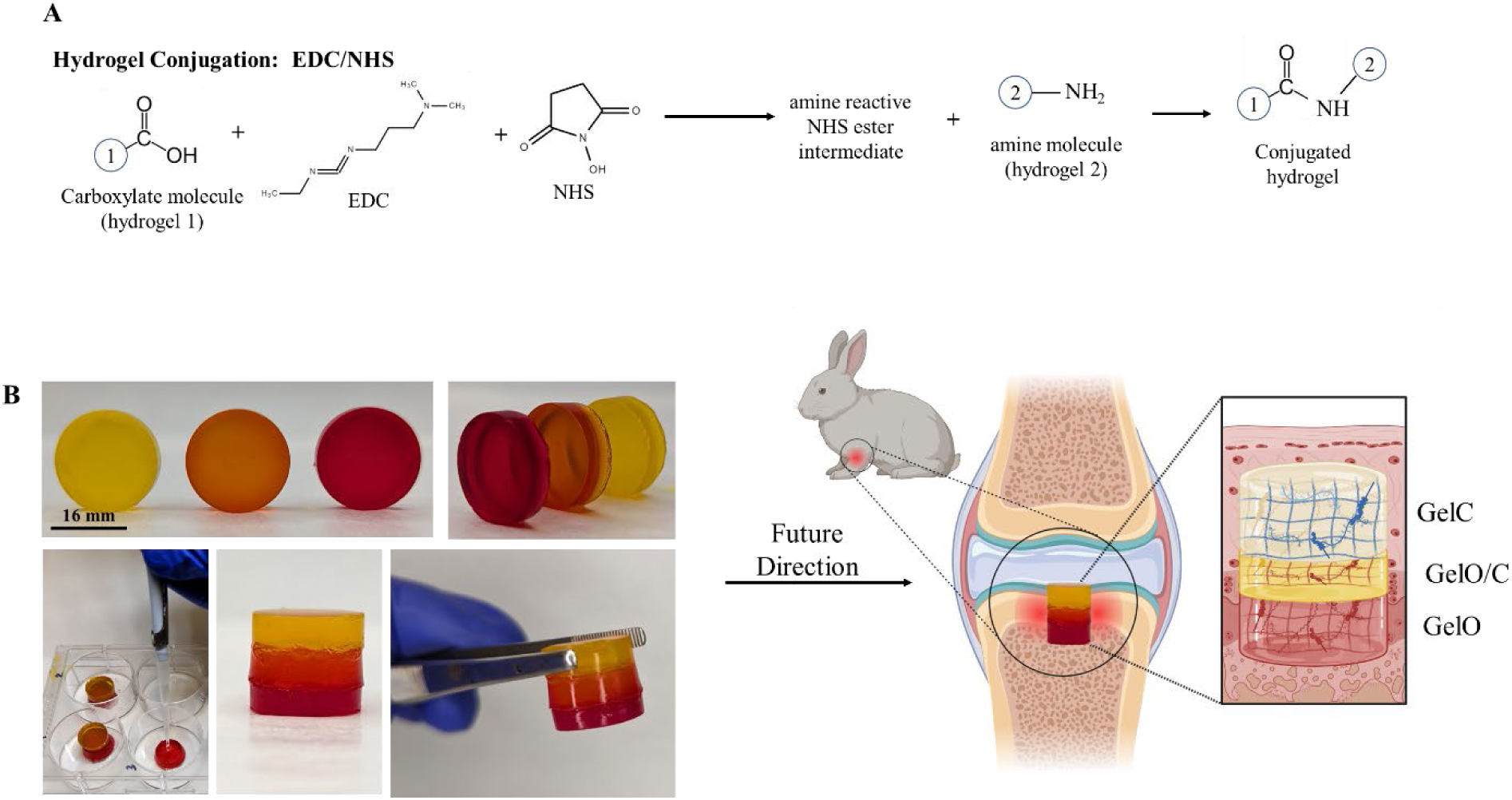
Conjugation of GelMA Hydrogels through EDC/NHS Coupling for Osteochondral Plug Design. A) The figure provides an illustration depicting the process of conjugating GelMA hydrogels. EDC (1-Ethyl-3-(3-dimethylaminopropyl) carbodiimide) and NHS (N-Hydroxysuccinimide) are used as coupling agents to covalently link the GelMA hydrogels together. This conjugation process facilitates the creation of a structurally stable and functional three-layer hydrogel. B) The figure presents a conceptual design for an osteochondral plug that can be potentially utilized in future applications. The osteochondral plug is composed of three distinct layers: a chondrogenic layer (GelC) to promote cartilage formation, an interphase layer (GelO/C) to facilitate the integration between cartilage and bone, and an osteogenic layer (GelO) to support bone regeneration. The GelMA hydrogels in each layer are conjugated through EDC/NHS coupling, ensuring structural integrity and compatibility. By implanting the three-layer hydrogel into the osteochondral defect site, the plug aims to facilitate the regeneration of both cartilage and bone, promoting functional tissue repair.

The conceptual design of the osteochondral plug, consisting of three distinct layers, showcases the versatility of the EDC/NHS coupled GelMA hydrogels in promoting osteochondral repair (Figure 9B). The first layer, the chondrogenic layer (GelC), specifically aims to stimulate cartilage formation. The second layer, the interphase layer (GelO/C), serves as a transitional zone between the cartilage and bone layers, which can facilitate a smooth and continuous surface for load transmission and joint function. The third layer, the osteogenic layer (GelO), focuses on promoting bone regeneration by providing a conducive environment for stem cell attachment and differentiation. By integrating the three-layer hydrogel into the osteochondral defect site, the developed osteochondral plug may have a promising effect on the regeneration of both cartilage and bone. Further investigations are needed to explore the materials’ mechanical properties, cell adhesion, overall biocompatibility, and differentiation potential.

## Conclusion

In this study, we developed a bioactive hydrogel-based scaffold for application in osteochondral tissue repair, combining dECM isolated from differentiated bone and cartilage cell lines with GelMA and DLP printing technology. The results of our experiments demonstrate the potential of this hydrogel as a drug-free and cell-free treatment option for osteoarthritis (OA) and small osteochondral defects (OCDs), simultaneously regenerating the articular cartilage and subchondral bone region of the osteochondral interface. The production and characterization of dECM from MC3T3-E1 and ATDC5 cell lines proved to be efficient and reproducible, yielding significant amounts of decellularized matrices with preserved bioactive molecules. SEM imaging and staining techniques confirmed the successful decellularization and provided insights into the structural and mineralization differences between differentiated and non-differentiated dECM. Furthermore, the quantitative analysis of calcium levels and sulfated glycosaminoglycans in the dECM supported their osteogenic and chondrogenic properties, respectively.

The physicochemical and mechanical characterization of GelO and GelC hydrogels revealed improved mechanical stiffness and reduced swelling and degradation profiles compared to the GelMA control group. These findings indicate that the incorporation of dECM positively influences the hydrogel’s mechanical properties, making it a more suitable candidate for osteochondral repair applications. Additionally, cytocompatibility studies demonstrated that both GelO and GelC hydrogels supported the survival, proliferation, and differentiation of hADSCs towards osteogenic and chondrogenic lineages. Alizarin Red and Alcian Blue staining confirmed the osteogenic and chondrogenic potential of GelO and GelC, respectively. Additionally, the combination of GelO and GelC in a bilayer configuration displayed a remarkable synergistic effect in promoting both osteogenesis and chondrogenesis, making it a promising candidate for osteochondral repair.

Furthermore, the covalent conjugation of GelMA hydrogels through EDC/NHS chemistry for the design of the osteochondral plug demonstrated its versatility in osteochondral scaffolding. The three-layer hydrogel, consisting of chondrogenic, interphase, and osteogenic layers, showcases the potential for regenerating both cartilage and bone in the osteochondral defect site.

While this study has provided significant insights into the development and characterization of the bioactive hydrogel, further investigations are warranted to explore its long-term performance, and *in vivo* biocompatibility. Additionally, optimization of the hydrogel composition, culture conditions, and material properties could further enhance the scaffold’s regenerative capabilities. Overall, this research contributes to the advancement of tissue engineering approaches for osteochondral tissue repair and brings us closer to providing effective and minimally invasive treatments for patients suffering from OA and OCDs.

## Acknowledgements

Arghya Paul is thankful for the funding and support from Canada Research Chairs Program of the Natural Sciences and Engineering Research Council (NSERC) of Canada (CRC-2018-00028), NSERC Discovery Grant, NSERC Discovery Accelerator Supplements (DAS), Early Research Award (ERA) from Province of Ontario, and Wolfe-Western Fellowship. Ali Coyle is grateful for the funding and support from NSERC Canada Graduate Scholarships—Masters program (CGS-M). Yasmeen Shamiya is thankful for the funding and support from the Ontario Graduate Scholarship (OGS). Aishik Chakraborty is supported in part by a Transdisciplinary Award from the Bone and Joint Institute, The University of Western Ontario, Canada. The authors would also like to acknowledge Mia Van Oirschot for her help with 3D printing hydrogels, Surface Science Western for scanning electron microscopy and energy dispersive x-ray, and SPAARC Biocentre at SickKids, Toronto for proteomics assay. The authors would also like to thank Prof. Subrata Chakrabarti and Prof. Kibret Mequanint for allowing access to their lab instruments, and Prof. Frank Beier for gifting ATDC5 cells. The authors are also thankful to the company BioRender. Some images and illustrations were created with **<u>BioRender.com</u>**.

## References

[1] D. K. Verma, P. Kumari, S. Kanagaraj, Annals of Biomedical Engineering 2022 50:3 2022, 50, 237.

[2] S. Davis, M. Roldo, G. Blunn, G. Tozzi, T. Roncada, Front Bioeng Biotechnol 2021, 9, 603408.

[3] S. P. Yu, D. J. Hunter, Aust Prescr 2015, 38, 115.

[4] R. Sommaggio, M. Uribe-Herranz, M. Marquina, C. Costa, Eur Cell Mater 2016, 32, 24.

[5] C. Vyas, · Hussein Mishbak, G. Cooper, C. Peach, R. F. Pereira, P. Bartolo, Biomanufacturing Reviews 2020 5:1 2020, 5, 1.

[6] E. Zeimaran, S. Pourshahrestani, A. Fathi, N. A. bin A. Razak, N. A. Kadri, A. Sheikhi, F. Baino, Acta Biomater 2021, 136, 1.

[7] D. E. Radulescu, I. A. Neacsu, A. M. Grumezescu, E. Andronescu, Polymers 2022, Vol. 14, Page 899 2022, 14, 899.

[8] C. H. Lee, A. Singla, Y. Lee, Int J Pharm 2001, 221, 1.

[9] E. C. Doyle, N. M. Wragg, S. L. Wilson, Knee Surg Sports Traumatol Arthrosc 2020, 28, 3827.

[10] D. Jovic, Y. Yu, D. Wang, K. Wang, H. Li, F. Xu, C. Liu, J. Liu, Y. Luo, Stem Cell Reviews and Reports 2022 18:5 2022, 18, 1525.

[11] X. Du, L. Cai, J. Xie, X. Zhou, Bone Res 2023, 11, DOI 10.1038/S41413-022-00239-4.

[12] S. Qin, J. Zhu, G. Zhang, Q. Sui, Y. Niu, W. Ye, G. Ma, H. Liu, Front Bioeng Biotechnol 2023, 11, 1127949.

[13] W. Wei, Y. Ma, X. Yao, W. Zhou, X. Wang, C. Li, J. Lin, Q. He, S. Leptihn, H. Ouyang, Bioact Mater 2021, 6, 998.

[14] D. C. Browe, P. J. Díaz-Payno, F. E. Freeman, R. Schipani, R. Burdis, D. P. Ahern, J. M. Nulty, S. Guler, L. D. Randall, C. T. Buckley, et al., Acta Biomater 2022, 143, 266.

[15] G. I. Barbulescu, F. M. Bojin, V. L. Ordodi, I. D. Goje, A. S. Barbulescu, V. Paunescu, International Journal of Molecular Sciences 2022, Vol. 23, Page 13040 2022, 23, 13040.

[16] Q. Yao, Y. W. Zheng, Q. H. Lan, L. Kou, H. L. Xu, Y. Z. Zhao, Materials Science and Engineering: C 2019, 104, 109942.

[17] N. Rajabi, A. Rezaei, M. Kharaziha, H. R. Bakhsheshi-Rad, H. Luo, S. Ramakrishna, F. Berto, Tissue Eng Part A 2021, 27, 679.

[18] A. Neishabouri, A. Soltani Khaboushan, F. Daghigh, A. M. Kajbafzadeh, M. Majidi Zolbin, Front Bioeng Biotechnol 2022, 10, 805299.

[19] A. Chakraborty, S. Pacelli, S. Alexander, S. Huayamares, Z. Rosenkrans, F. E. Vergel, Y. Wu, A. Chakravorty, A. Paul, Mol Pharm 2022, DOI 10.1021/ACS.MOLPHARMACEUT.2C00564/ASSET/IMAGES/LARGE/MP2C00564_0004.JPEG.

[20] B. J. Klotz, D. Gawlitta, A. J. W. P. Rosenberg, J. Malda, F. P. W. Melchels, Trends Biotechnol 2016, 34, 394.

[21] S. Krishnamoorthy, B. Noorani, C. Xu, Int J Mol Sci 2019, 20, DOI 10.3390/IJMS20205061.

[22] C. E. Choi, A. Chakraborty, H. Adzija, Y. Shamiya, K. Hijazi, A. Coyle, A. Rizkalla, D. W. Holdsworth, A. Paul, Gels 2023, 9, 923.

[23] J. Li, A. D. Celiz, J. Yang, Q. Yang, I. Wamala, W. Whyte, B. R. Seo, N. V. Vasilyev, J. J. Vlassak, Z. Suo, et al., Science 2017, 357, 378.

[24] C. A. Gregory, W. G. Gunn, A. Peister, D. J. Prockop, Anal Biochem 2004, 329, 77.

[25] M. W. Pfaffl, Nucleic Acids Res 2001, 29, E45.

[26] M. Izumiya, M. Haniu, K. Ueda, H. Ishida, C. Ma, H. Ideta, A. Sobajima, K. Ueshiba, T. Uemura, N. Saito, et al., Int J Mol Sci 2021, 22, DOI 10.3390/IJMS22147752/S1.

[27] Y. Yao, Y. Wang, J Cell Biochem 2013, 114, 1223.

[28] D. Wilhelm, H. Kempf, A. Bianchi, J. B. Vincourt, J Proteomics 2020, 219, 103718.

[29] M. Yuasa, T. Yamada, T. Taniyama, T. Masaoka, W. Xuetao, T. Yoshii, M. Horie, H. Yasuda, T. Uemura, A. Okawa, et al., PLoS One 2015, 10, DOI 10.1371/JOURNAL.PONE.0116462.

[30] X. Liu, J. Liu, N. Kang, L. Yan, Q. Wang, X. Fu, Y. Zhang, R. Xiao, Y. Cao, Int J Mol Sci 2014, 15, 1525.

[31] K. Fujisawa, K. Hara, T. Takami, S. Okada, T. Matsumoto, N. Yamamoto, I. Sakaida, Stem Cell Res Ther 2018, 9, 1.

[32] F. Langenbach, J. Handschel, Stem Cell Res Ther 2013, 4, 1.

[33] I. Alesutan, F. Moritz, T. Haider, S. Shouxuan, C. Gollmann-Tepeköylü, J. Holfeld, B. Pieske, F. Lang, K. U. Eckardt, S. S. Heinzmann, et al., J Mol Med (Berl) 2020, 98, 985.

[34] W. Weng, F. Zanetti, D. Bovard, B. Braun, S. Ehnert, T. Uynuk-Ool, T. Histing, J. Hoeng, A. K. Nussler, R. H. Aspera-Werz, J Mater Sci Mater Med 2021, 32, 1.

[35] N. A. M. Bax, M. H. van Marion, B. Shah, M. J. Goumans, C. V. C. Bouten, D. W. J. Van Der Schaft, J Mol Cell Cardiol 2012, 53, 497.

[36] Y. K. Kim, L. S. Gu, T. E. Bryan, J. R. Kim, L. Chen, Y. Liu, J. C. Yoon, L. Breschi, D. H. Pashley, F. R. Tay, Biomaterials 2010, 31, 6618.

[37] M. Koblenzer, M. Weiler, A. Fragoulis, S. Rütten, T. Pufe, H. Jahr, Cells 2022, 11, 2702.

[38] J. Chen, K. Singh, B. B. Mukherjee, J. Sodek, Matrix 1993, 13, 113.

[39] M. Ikegame, S. Ejiri, H. Okamura, Journal of Histochemistry and Cytochemistry 2019, 67, 107.

[40] S. Kim, S. S. Lee, B. Son, J. A. Kim, N. S. Hwang, T. H. Park, ACS Biomater Sci Eng 2021, 7, 1134.

[41] G. Chen, C. Deng, Y. P. Li, Int J Biol Sci 2012, 8, 272.

[42] G. K. Hunter, D. P. Holmyard, K. P. H. Pritzker, J Cell Sci 1993, 104 (Pt 4), 1031.

[43] D. Magne, G. Bluteau, C. Faucheux, G. Palmer, C. Vignes-Colombeix, P. Pilet, T. Rouillon, J. Caverzasio, P. Weiss, G. Daculsi, et al., Journal of Bone and Mineral Research 2003, 18, 1430.

[44] C. A. Paggi, J. Hendriks, M. Karperien, S. Le Gac, Lab Chip 2022, 22, 1815.

[45] V. Lefebvre, M. Dvir-Ginzberg, Connect Tissue Res 2017, 58, 2.

[46] M. Strecanska, L. Danisovic, S. Ziaran, M. Cehakova, Life 2022, Vol. 12, Page 2066 2022, 12, 2066.

[47] X. Wang, Y. Guan, S. Xiang, K. L. Clark, P. G. Alexander, L. E. Simonian, Y. Deng, H. Lin, Front Cell Dev Biol 2022, 10, 812081.

[48] J. Cui, J. Zhang, International Journal of Molecular Sciences 2022, Vol. 23, Page 9253 2022, 23, 9253.

[49] N. Alcorta-Sevillano, I. Macías, A. Infante, C. I. Rodríguez, Cells 2020, Vol. 9, *Page* 2630 2020, 9, 2630.

[50] H. Li, J. Dai, Z. Wang, H. Zheng, W. Li, M. Wang, F. Cheng, Aggregate 2023, 4, e270.

[51] D. Ege, V. Hasirci, ACS Appl Bio Mater 2023, DOI 10.1021/ACSABM.3C00093/ASSET/IMAGES/LARGE/MT3C00093_0009.JPEG.

[52] Y. Piao, H. You, T. Xu, H. P. Bei, I. Z. Piwko, Y. Y. Kwan, X. Zhao, Engineered Regeneration 2021, 2, 47.

[53] T. Grossner, U. Haberkorn, J. Hofmann, T. Gotterbarm, International Journal of Molecular Sciences 2022, Vol. 23, Page 6288 2022, 23, 6288.

[54] Y. Jung, S. H. Kim, S. H. Kim, Y. H. Kim, J. W. Rhie, S. H. Kim, Macromol Res 2012, 20, 709.

[55] M. A. Aronow, L. C. Gerstenfeld, T. A. Owen, M. S. Tassinari, G. S. Stein, J. B. Lian, J Cell Physiol 1990, 143, 213.

[56] J. H. Xu, Z. H. Li, Y. D. Hou, W. J. Fang, Am J Transl Res 2015, 7, 2527.

[57] H. Chen, F. Y. Ghori-Javed, H. Rashid, M. D. Adhami, R. Serra, S. E. Gutierrez, A. Javed, J Bone Miner Res 2014, 29, 2653.

[58] J. Liao, T. Tian, S. Shi, X. Xie, Q. Ma, G. Li, Y. Lin, Bone Research 2017 5:1 2017, 5, 1.

[59] P. Comeau, T. Willett, P. Comeau, T. Willett, Macromol Mater Eng 2021, 306, 2000604.

[60] A. J. Kuijpers, G. H. M. Engbers, J. Krijgsveld, S. A. J. Zaat, J. Dankert, J. Feijen, 10.1163/156856200743670 2012, 11, 225.

[61] H. Goodarzi, K. Jadidi, S. Pourmotabed, E. Sharifi, H. Aghamollaei, Int J Biol Macromol 2019, 126, 620.

[62] N. Yin, Y. Han, H. Xu, Y. Gao, T. Yi, J. Yao, L. Dong, D. Cheng, Z. Chen, Materials Science and Engineering: C 2016, 59, 958.

[63] C. Lee, J. Shin, J. S. Lee, E. Byun, J. H. Ryu, S. H. Um, D. I. Kim, H. Lee, S. W. Cho, Biomacromolecules 2013, 14, 2004.

